# Glucocorticoids Modulate mRNA Translation Fate Through P-Body Dynamics

**DOI:** 10.1101/2025.10.30.685488

**Authors:** Victoria J. Nicolini, Mathilde Dupart, Wolfgang Raffelsberger, Olivia Vidal-Cruchez, Tifenn Rete, Karine Jacquet, Frédéric Brau, Sophie Abelanet, Marie Irondelle, Antonin Bourdin, Jérôme Durivault, Célia Gotorbe, Adeline Knittel-Obrecht, Bruno Didier, Pascal Villa, Christophe Di Giorgio, Baharia Mograbi, Arnaud Hubstenberger, Paul Hofman, Patrick Brest

**Author notes:** Correspondence to Patrick Brest. Université Côte d’Azur, IRCAN, 28 avenue de Valombrose, 06107 Nice, France.

## Abstract

Processing bodies (P-bodies) are cytoplasmic, membraneless organelles that play a key role in regulating RNA translation. To identify new pathways controlling their formation, we conducted a Food and Drug Administration (FDA)-approved drug screen. We found that glucocorticoids, among the most prescribed medicines, significantly increase P-body numbers across diverse epithelial cell types. This effect was fully reversible after glucocorticoid withdrawal, illustrating the adaptive dynamics of P-bodies. Using genetic invalidation and rescue approaches, we demonstrated that this accumulation requires the Glucocorticoid Receptor alpha isoform. P-body accumulation was associated with the sequestration of P-body-specific-targeted mRNAs, altering their translation yield. Notably, this translational regulation depends on transcript sequence features rather than abundance, with AU-rich mRNA transcripts being sequestered and GC-rich mRNAs preferentially translated under glucocorticoid treatment. Furthermore, we linked the decrease of LSM14B, a negative regulator of P-bodies, under glucocorticoid treatment to P-body reshaping. Our results reveal that, beyond their known transcriptional activity, prolonged exposure to glucocorticoids influences mRNA post-transcription and translation through a nucleotide composition-based mechanism.

## Results and Discussion

Processing-bodies (P-bodies) are cytoplasmic membraneless organelles that regulate mRNA translation through phase separation^1^. Despite extensive molecular characterization, the physiological signals controlling their assembly remain poorly understood. Sodium arsenite, one of the most used P-body chemical regulator, cause rapid cell death, limiting insights into specific regulatory mechanisms^1^.

Through an unbiased screen of 1,520 FDA-approved drugs, we identified glucocorticoids as potent inducers of P-body formation, as illustrated in Figure 1A (and in the supplementary data), establishing a previously unknown connection between steroid hormone signaling and cytoplasmic RNA granule dynamics. Glucocorticoids such as dexamethasone and prednisolone, among the most prescribed medication worldwide, increased P-body numbers at clinically relevant concentrations (0.1-1 μM) (Fig. 1B) within 48 hours across multiple cell lines (A549, HeLa, and Mel501) (Fig. 1C). To assess GR involvement and system dynamics, we used complementary approaches: GR antagonism (RU486) completely blocked P-body formation and FKBP5 upregulation, while glucocorticoid withdrawal rapidly reversed the phenotype within 24 hours (Fig. 1D-E). This GR-dependent, reversible response defines a tightly regulated pathway.

**Figure 1:**
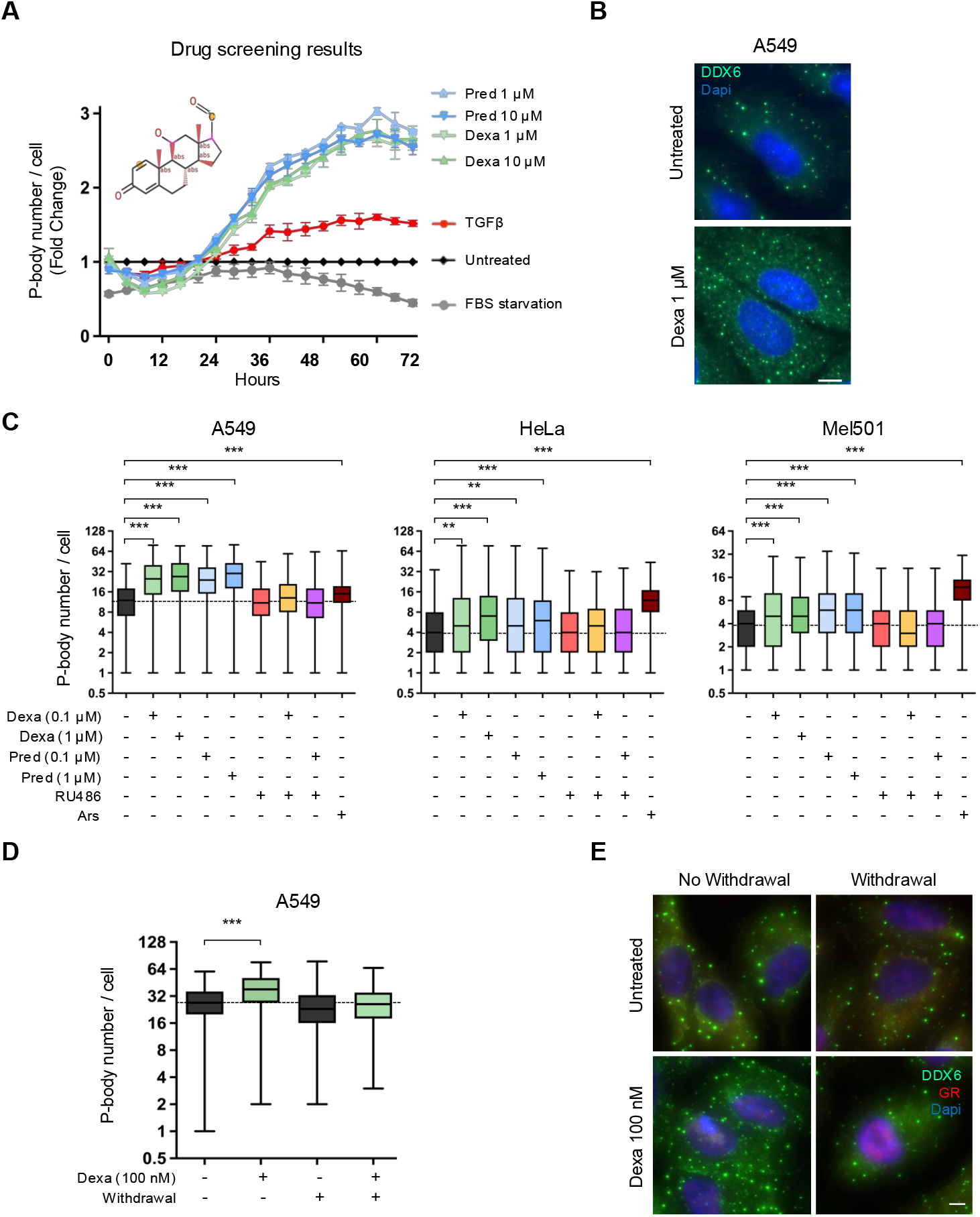
Glucocorticoid receptor activation increases P-body numbers in different epithelial cell lines. **(A)** Results of the drug screening showing the fold change ratio of P-body number per cell in A59 GFP-DDX6 cells treated with either 1 or 10 μM of dexamethasone (Dexa) or prednisolone (Pred) respectively, compared to control (0.1% DMSO). TGFb (10 ng/ml) and serum starvation served as a positive control for P-body induction and disassembly, respectively. A schematic representation shows the conserved molecular scaffold identified from the glucocorticoid screen; the orange dot indicates the position of structural variation (single or double bond in the six-membered ring, with potential carbonyl substitution) tolerated for P-body induction activity. **(B)** Representative immunofluorescence images of P-body marker (DDX6, green) and nuclei (Dapi, blue) in A549 cells treated with 1 μM Dexa for 48h. **(C)** Quantification of P-body number per cell in three different cell lines (A549, HeLa and Mel501) treated for 48 hours with Dexa at 0.1 or 1 μM, Pred at 0.1 or 1 μM, RU486 at 1μM (a glucocorticoid inhibitor, 1h pre-treatment), or sodium arsenite (Ars) at 0.5 mM for 30 minutes (a positive control for P-body induction). **(D)** Quantification of P-body number per cell in A549 cells treated with 100 nM of Dexa, after 48h of treatment medium was replaced with normal medium for 24h (Withdrawal) and **(E)** its corresponding representative immunofluorescence images with P-body marker (DDX6, green), glucocorticoid receptor (GR, red), and nuclei (Dapi, blue) staining. Scale bars: 10 μm. P-bodies were quantified in at least 100 cells per condition from at least three independent biological experiments. After normality testing (Shapiro’s test), data were analyzed using Kruskal-Wallis test with Dunn’s post-hoc comparison. Significance: p < 0.05 (* p < 0.05, ** p < 0.01, *** p < 0.001, ns = non-significant).

Using genetic invalidation (Fig. 2A-B) and rescue approaches (Fig. 2C-D), we demonstrated that the GRα isoform specifically, but not GRβ or truncated variants^3^, was both necessary and sufficient for P-body induction. Dose-response studies revealed an inverted U-shaped curve with GRα overexpression (Fig. 2E), indicating that both insufficient and excessive GR signaling can disrupt optimal P-body assembly, suggesting precise cellular calibration of this pathway. Importantly, glucocorticoid-induced P-bodies form independently of G3BP1-positive stress granules, distinguishing this response from classical stress responses^1^.

**Figure 2:**
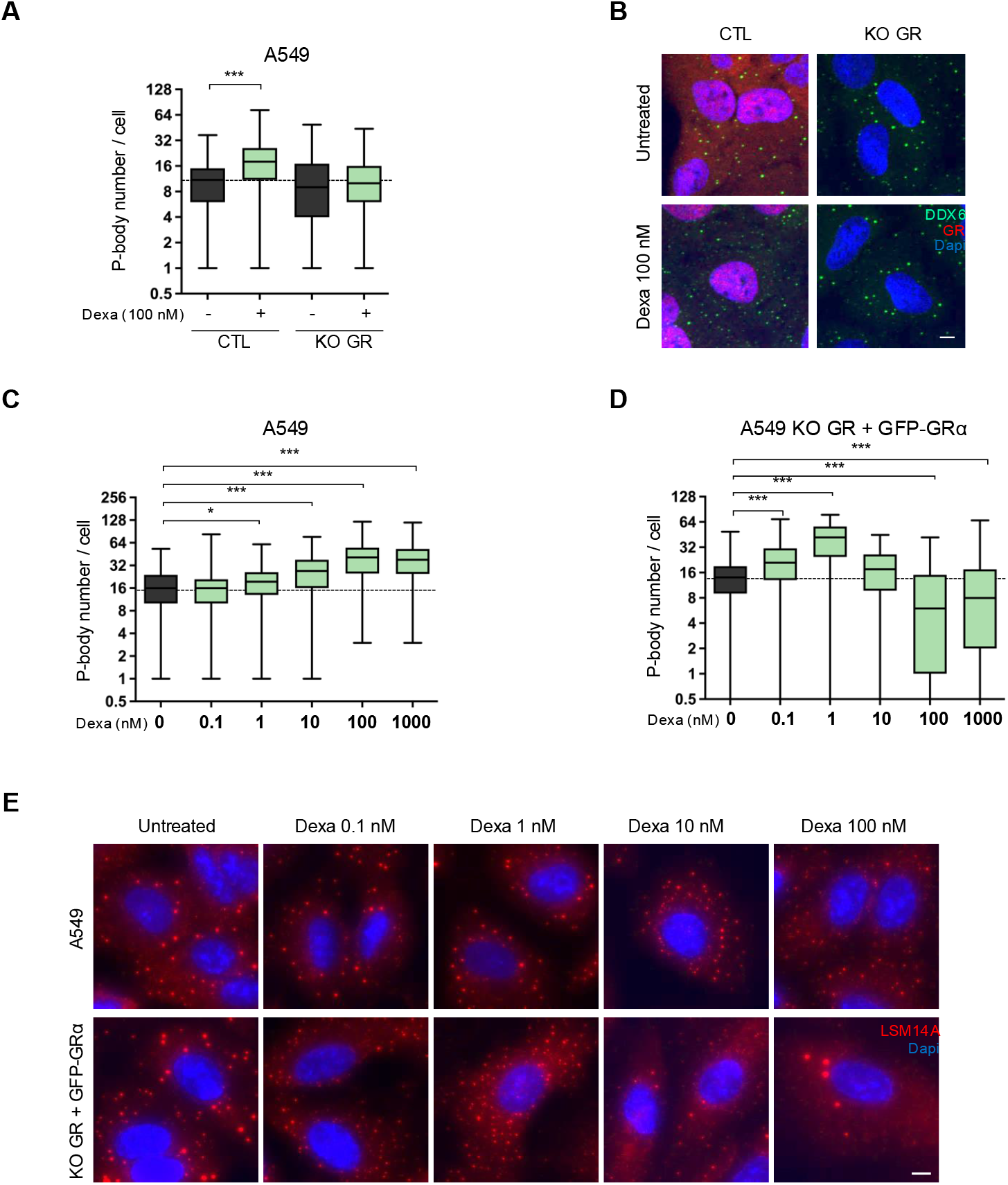
Glucocorticoid receptor alpha isoform is required for P-body increased number under glucocorticoid treatment. **(A)** Quantification of P-body number per cell in A549 control cells (CTL) or GR knockout (CRISPR GR KO) cells treated with 100 nM Dexa for 48 hours at and **(B)** its corresponding representative immunofluorescence images of DDX6 (P-body marker, green), GR (red) and the nuclei (Dapi, blue) from the same conditions. **(C-D)** Quantification of P-body number per cell in **(C)** A549 CTL and **(D)** A549 KO GR + GFP-GRa cells treated from 48 hours with Dexa ranging from 0.1 nM to 1 μM and **(E)** its corresponding representative immunofluorescence images of P-body marker (LSM14A, red) and nuclei (Dapi, blue) with Dexa ranging from 0.1 to 100 nM. Scale bars: 10 μm. P-bodies were quantified in at least 100 cells per condition from at least three independent biological experiments. After normality testing (Shapiro’s test), data were analyzed using Kruskal-Wallis test with Dunn’s post-hoc comparison. Significance: p < 0.05 (* p < 0.05, ** p < 0.01, *** p < 0.001, ns = non-significant).

Unlike translation inhibitors such as cycloheximide, we demonstrate that glucocorticoids do not induce a general shutdown of protein production (Fig. 3A). Indeed, SUnSET assay^4^ showed sustained protein synthesis under glucocorticoid treatments, indicating a selective rather than widespread response. To understand this selectivity, we conducted parallel transcriptomics and proteomics after GR activation (Fig. 3B-C). Both datasets exhibited robust glucocorticoid gene enrichment signature (GSEA NES = 2.54 and 2.71, p<10^-10^ respectively). However, by analyzing how GR activation modifies differential mRNA-protein correlations, we revealed a striking selectivity: while canonical GR(*NR3C1*)-induced gene signatures are correlated (Fig. 3D), P-body-targeted transcripts exhibited remarkable decorrelation with mRNA accumulation (NES = 1.92, p < 10^-10^) (Fig. 3E) but protein depletion (NES = −1.84, p < 2.1 10^-6^), in agreement with previous data^5^. This selective P-body-targeted transcripts accumulation proved to be sequence-dependent, as previously published^6^. Indeed, these gene sequences displayed a significant AU-rich composition bias in total, CDS, and 3’UTR regions in comparison with whole mRNA or canonical GR-induced gene sequences. Furthermore, by analyzing genes with lower-than-expected protein levels genome-wide (Fig. 3F-G), we confirmed that this nucleotide composition-based selectivity found in purified P-bodies expended at the cellular-wide level under GR activation affecting substantially more transcripts with AU-rich composition. Reciprocally, genes with GC-rich composition in total, CDS, and 3’UTR regions displayed higher-than-expected protein levels. Overall, AU/GC-induced translational selectivity extends beyond P-body-associated transcripts under glucocorticoid treatment.

**Figure 3:**
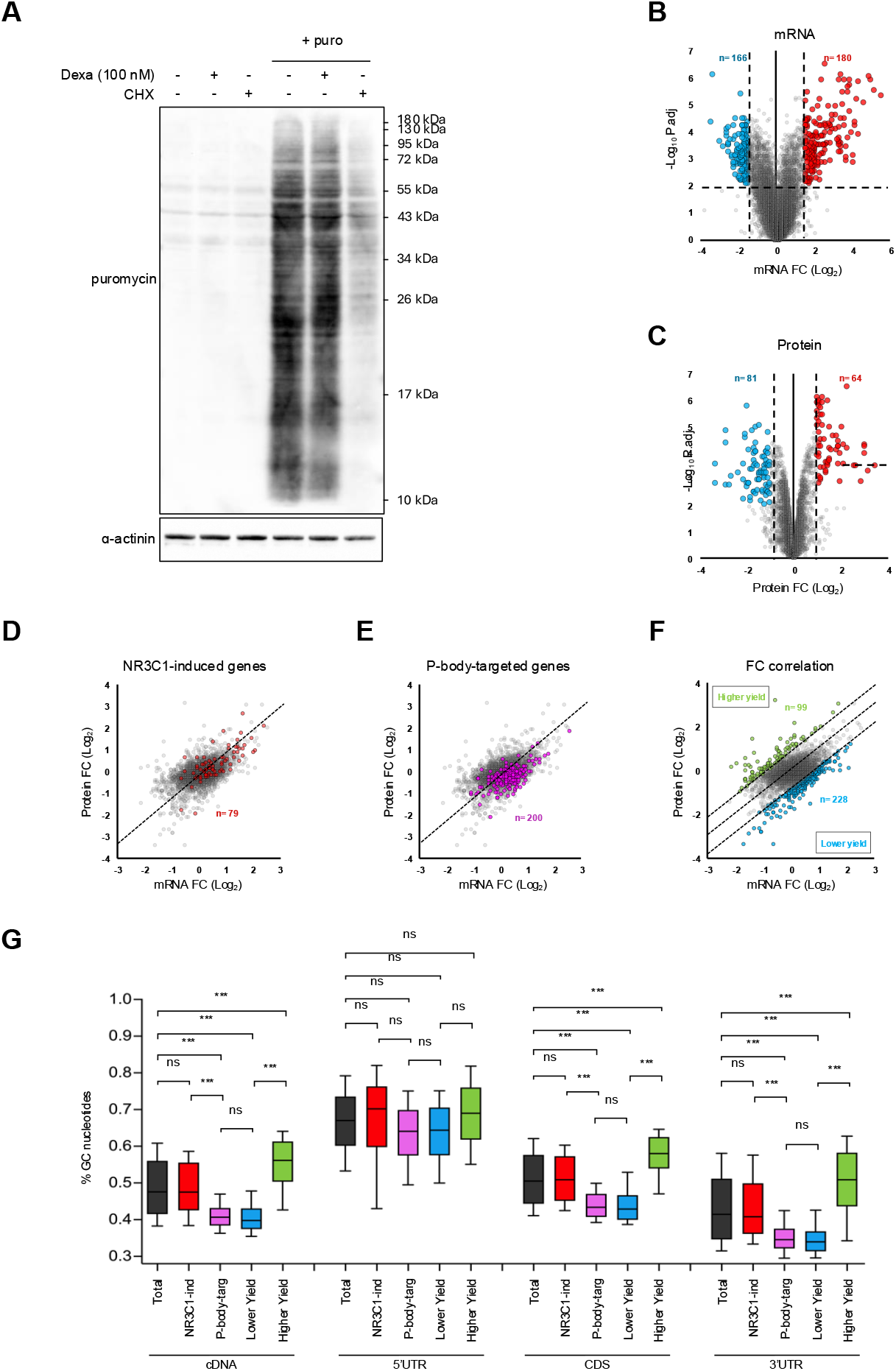
GR activation promotes a P-bodies-related AU-rich bias in mRNA translation fate. **(A)** Translation activity through SUnSET assay was assessed by immunoblot analysis with anti-puromycin antibodies (and anti-a-actinin antibodies were used as loading control). A549 cells were treated with 100 nM of Dexa for 48 hours, labeled or not with puromycin (puro) at 10 μg/ml for 10 minutes, in presence or absence of cycloheximide (CHX, translation inhibitor) at 30 μg/ml for 1 hour. **(B-C)** Volcano plots depicting differential **(B)** mRNA (ribo-depleted RNAseq) or **(C)** protein (mass-spectrometry) expression in dexamethasone-treated versus untreated A549 cells. Red dots indicate significantly upregulated genes (fold change >1.5, p <0.01 for mRNA and fold change > 1.2, p <0.01 for proteins respectively), while blue dots represent significantly downregulated transcripts. Statistical significance (−Log10 p-value adjusted) is plotted against magnitude of change (Log2 fold change). **(D-G)** Fold change values for transcripts and proteins following 48 h dexa treatment were plotted on the same graph to assess the correlation between mRNA and protein level changes. Specific genes signatures were highlighted: **(D)** NR3C1-induced genes (NR3C1-ind, red), **(E)** P-body-targeted genes (P-body-targ, purple), and **(F)** genes stratified by protein/mRNA ratio: those with elevated protein relative to mRNA (green) and those with reduced protein relative to mRNA (blue). **(G)** GC content of each gene category (extracted from Ensembl) across mRNA regions (5’UTR, CDS, 3’UTR, full-length cDNA), stratified by the gene signatures shown above. After normality testing (Shapiro’s test), data were analyzed using Kruskal-Wallis test with Dunn’s post-hoc comparison. Significance: p < 0.05 (* p < 0.05, ** p < 0.01, *** p < 0.001, ns = non-significant).

Restricted proteomic analysis of P-body proteins under glucocorticoid treatment revealed LSM14B downregulation as a possible key downstream regulator of this GRα-dependent P-body reshaping (Fig. 4A). Indeed, after dexamethasone treatment, the number of LSM14B-positive P-bodies decreased significantly. Interestingly, while its paralogue LSM14A is a core essential component of P-bodies (as DDX6 and 4-ET) and acts as a translational repressor^7^, LSM14B controls oocyte mRNA storage and stability to ensure female fertility^8^. Likewise glucocorticoid treatment, siRNA-mediated knockdown of LSM14B recapitulated the GRα-induced P-body phenotype (Fig. 4B-C), while overexpressing LSM14B prevented it (Fig. 4D-E), demonstrating causality. Interestingly the number of LSM14B-positive P-bodies mirrored the previously observed dynamic (Fig. 1D-E) returning to baseline number within 24 hours of glucocorticoid withdrawal (Fig. 4F).

**Figure 4:**
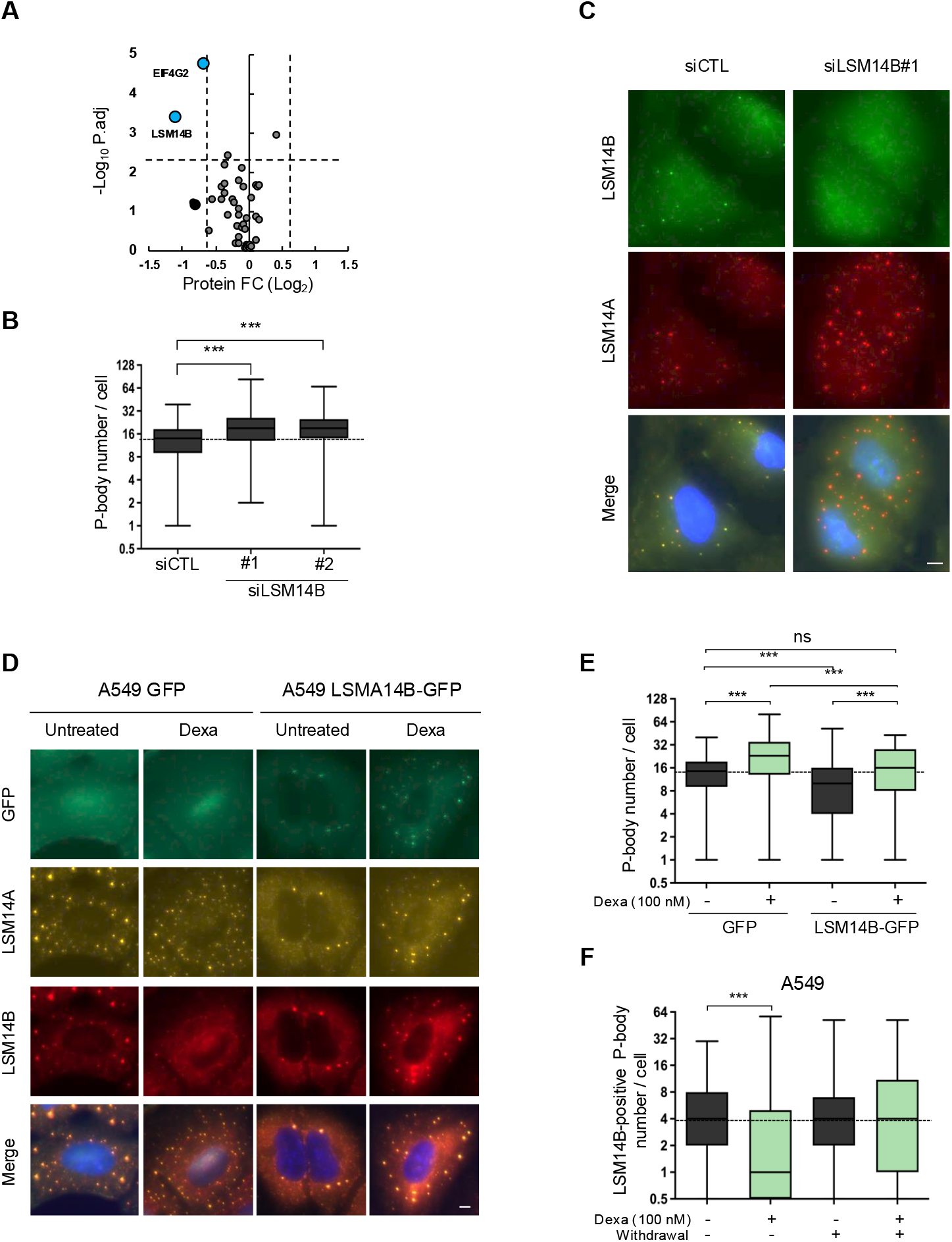
LSM14B expression is inversely correlated with GR-dependent P-body increase. **(A)** Volcano plot depicting P-body-specific differential protein expression in A549 cells treated with dexamethasone versus untreated controls. Blue dots indicate significantly downregulated proteins (fold change >-0.6, p <0.01). Statistical significance (−Log10 p-value adjusted) is plotted against magnitude of change (Log2 fold change). **(B)** Quantification of P-body number per cell in A549 cells transfected with siCTL or two different siRNAs against LSM14B (siLSM14B#1/#2) and **(C)** its corresponding representative immunofluorescence images with LSM14B (green), P-body marker (LSM14A, red), and nuclei (Dapi, blue) stainings. **(D)** Immunofluorescence images of GFP constructs (green), P-body marker (LSM14A, yellow), LSM14B (red), and nuclei (Dapi, blue) staining in A549 GFP and A549 LSM14B-GFP cells treated with 100nM of dexa for 48 hours and **(E)** its quantification of P-body number per cell. **(F)** P-body numbers quantified in A549 cells treated with 100 nM dexamethasone for 48 hours, then cultured in normal growth conditions for 24 hours (withdrawal). Scale bars: 10 μm. P-bodies were quantified in at least 100 cells per condition from at least three independent biological experiments. After normality testing (Shapiro’s test), data were analyzed using Kruskal-Wallis test with Dunn’s post-hoc comparison. Significance: p < 0.05 (* p < 0.05, ** p < 0.01, *** p < 0.001, ns = non-significant).

To investigate the mechanism underlying glucocorticoid-induced LSM14B downregulation, we conducted a systematic analysis that revealed post-transcriptional regulation since RNA-seq showed increased mRNA levels concurrent with decreased protein expression following GR alpha activation. In this context, we systematically excluded several regulatory pathways: GR depletion did not affect baseline LSM14B levels, ruling out direct cytoplasmic GR-mRNA interactions; LSM14A levels remained unchanged, excluding indirect regulation through its paralog; LSM14B protein is stable under basal conditions after cycloheximide treatment demonstrating absence of post-translational regulation. Interestingly, differential responses between endogenous and exogenous LSM14B seem to implicate 3’UTR-dependent regulation (supplementary data). However, no canonical GR-responsive RBP or microRNA binding sites were identified in the LSM14B 3’UTR highlighting possible non-canonical binding mechanisms. Although the precise molecular mediators remain to be identified, our complementary gain- and loss-of-function experiments unequivocally establish the functional link between GR signaling and LSM14B regulation, providing a framework for future mechanistic studies.

The marked increase in P-body numbers amplifies selective AU-rich mRNA accumulation cellular-wide, transforming a localized regulatory organelle into genome-wide reshaping of translation patterns. The mRNA content-dependent partitioning mechanism provides a conceptual framework that extends to stress contexts. AU-rich transcripts are preferentially sequestered, while GC-rich housekeeping transcripts remain available for translation, allowing glucocorticoids to regulate cellular proteomes without direct transcriptional control. As essential stress hormones coordinating complex adaptive responses, glucocorticoids may use P-body remodeling as a conserved mechanism for rapid proteomic reprogramming during stress adaptation. The preferential sequestration of AU-rich mRNAs is particularly significant, as these transcripts predominantly encode inflammatory mediators, immune regulatory factors, and RNA-binding proteins^9^. This selective sequestration aligns mechanistically with the well-established anti-inflammatory effects of glucocorticoids, suggesting that P-body remodeling represents a conserved additive strategy for suppressing inflammatory gene expression.

These findings expand our understanding of glucocorticoid action beyond the canonical transcriptional model, revealing an unexpected intersection between hormonal signaling and RNA translation fate. After the well-known glucocorticoid associated transcriptional response, this P-body associated post-transcriptional regulation is particularly relevant for chronic glucocorticoid exposure, which commonly occurs in clinical settings and can possibly lead to Cushing syndrome and other complications. The “stealthy” nature of this delayed response may explain why side effects emerge insidiously during prolonged therapy. Notably, reversibility upon withdrawal in our cell-based system suggests that some glucocorticoid-induced effects may be partly reversible, opening possibilities for sequential dosing regimens to possibly prevent pathological P-body accumulation and associated adverse glucocorticoid complications^3^.

In conclusion, this work reveals a novel post-transcriptional dimension of glucocorticoid action, expanding our understanding of how these essential stress hormones and widely used therapeutic agents exert their complex physiological and clinical effects.

## Materials and Methods

### Cell culture

A549 (human lung adenocarcinoma epithelial cell line, LUAD, ATCC Cat# CCL-185, RRID: CVCL_0023), HeLa (human uterus; cervix adenocarcinoma epithelial cell line, ATCC Cat# CCL-2, RRID: CVCL_0030), and Mel501 (melanoma epithelial cell line, SKCM, RRID: CVCL_4633, kind gift from Dr. Gillot Laboratory) cells were cultured according to the recommendations of the ATCC. A549 and HeLa cells were grown in Dulbecco’s modified eagle medium (DMEM) while Mel501 cells were cultured in RPMI 1640, both supplemented with 5% fetal bovine serum (FBS). All cells were maintained at 37°C in a 5% CO_2_ humidified incubator. Cells were cultured for no more than one month and were identified using STR profiling (Eurofins Genomics, Ebersberg, Germany). Mycoplasma contamination was checked every two weeks by Plasmotest (Invivogen, France).

### Treatments

For experimental treatments, cells were exposed to TGFβ (Sigma, Cat# SRP3171) at 10 ng/ml for 72h, two different glucocorticoids: dexamethasone (Bertin Bioreagent, Cat# 20340-100) or prednisolone (Bertin Bioreagent, Cat# 15933-1) at varying concentrations and varying time, RU486 (Cayman Chemical, Cat# 10006317) at 1 μM for 1h pre-treatment, sodium arsenite (NaAsO_2_) (Sigma, Cat#1.06771000) at 0.5 mM for 30 min, cycloheximide (CHX) (Sigma Cat#C6255) at 30 μg/ml for 1h and puromycin (puro) (Inivogen, Cat# ant-pr-1) at 10 μg/ml for 10 min.

Cells were plated at a density of 200,000 cells in 6-well plates. After 24h, cells were transfected with either a negative control siRNA (Silencer Select® small interfering RNA) (Thermo Fisher Cat#4390844) or with pre-validated LSM14B siRNA (s229767, s45634, Thermo Fisher) using JetPrime (PolyPlus) according to the manufacturer’s instructions. Cells were harvested by scraping 48h after transfection for further analysis.

### Drug screening

A high-throughput drug high-content screening was performed on a fully automated robotic platform (Plateforme de Chimie Biologique Intégrative de Strasbourg (PCBIS) using the Prestwick Chemical Library containing 1,520 FDA-approved compounds. A549 cells stably expressing DDX6-GFP, a marker of P-bodies, were seeded in 96-well plates (Greiner) and treated with compounds at 10 μM for 72h with a final DMSO (compound solvent) concentration of 0.1%, applied across all conditions, including controls. Images (phase contrast and fluorescence for GFP) were acquired every 4h using the IncuCyte-S3® system (Sartorius) and then analyzed with the dedicated Incucyte software to have access to the cell confluency and the P-body number per cells. Compounds were selected based on two stringent criteria: cell confluency at 72h (minimum 65%) to exclude cytotoxic effects, and P-body numbers exceeding TGFβ induction. 97 promising hits (6.4%) were then subjected to secondary validation assays at both 1 μM and 10 μM to confirm their effects and assess dose dependency.

CRISPR-Cas9 gene editing A549 wildtype (WT) cells were transfected using jetPEI (Polyplus, 101000053) according to the manufacturer’s instructions with the pSpCas9(BB)-2A-GFP (PX458) plasmid (a gift from Feng Zhang; Addgene plasmid #48138; RRID: Addgene_48138) containing clustered regularly interspaced short palindromic repeats (CRISPR)-CRISPR-associated protein 9 (Cas9) targeting the following regions : exon 2 (5’-GTAGAAAAAACTGTTCGACCA-3’), exon3 (5’-GAGCTCCTCAACAGCAACAAC-3’), and exon 5 (5’-GAACCTCCAACAGTGACACCA-3’and 5’-GCGCTCAACATGTTAGGAGGG-3’) targeting the NR3C1 gene. GFP-positive cells were single-cell sorted by flow cytometry (BD FACSMelody) 24h post-transfection into 96-well plates containing DMEM with 10% FBS. NR3C1 knockout (KO GR cells) clones were screened by Western blotting and validated by Sanger sequencing (Eurofins Genomics). Two independent knockout clones were selected for experiments to minimize clonal effects.

### Plasmid Constructs

The pPRIPu GFP-DDX6 plasmid used in this study was constructed as follows: the pPRIPu CrUCCI vector (kind gift from Dr. Delaunay) was amplified with primer adaptors containing AgeI and BamHI restriction sites. pEGFP-C1_p54cp plasmid (kind gift from Drs. Weil and Kress) was digested by AgeI and BamHI, and the obtained GFP-DDX6 fragment was inserted in pPRIPU after digestion with AgeI and BamHI. The pPRIPu GFP-GR plasmids used in this study were constructed as follows: the eGFP-NR3C1 (GRα) insert was digested with AgeI and BamHI from pEGFP GR (Addgene Plasmid #47504) and inserted into pPRIPU after digestion with AgeI and BamHI. GRβ, GRΔEx9 and GRα-RK491AA isoforms were generated from pPRIPu GFP-GRα by reverse PCR using specific primers (GRβ_Fwd: 5’-AGCACATCTCACACATTAATCTGAGGATCCACCGGATCTAGATAACTG-3’; GRβ_Rev: 5’-TTCTGGTTTTAACCACATAACATTTTCATGCATAGAATCCAAGAGTTTTGTCAG-3’; GRΔEx9_Fwd: 5’-TGAGGATCCACCGGATCTAGATAACTG-3’; GRΔEx9_Rev: 5’-TTCATGCATAGAATCCAAGAGTTTTGTCAG-3’). The pPRIPu GFP-LSM14B plasmid used in this study was constructed as follows: LSM14B cDNA was generated using a 5’-CTTTGGCTGCACCCTCACAC-3’ primer using SuperscriptIV enzyme (Invitrogen). Then the LSM14B cDNA was amplified with primer adaptors containing XhoI and BglII restriction sites (LSM14B_Fwd: 5’-GCCCCGGTAGATCTCGGGCCCGCGGTACCGTCGAGTCACACCCTGCCAGTCCC-3’ and LSM14B_Rev: 5’-CAGATCTCGAGCTCAAGCTTCGAATTCCATGAGCGGCTCCTCAGG-3’) and inserted into pPRIPu GFP-DDX6 previously digested by XhoI and BamHI (compatible with BglII). For all plasmids, the integrity of the entire sequence was confirmed by sequencing (Eurofins Genomics, Snapgene).

### Generation of stable cell lines

Briefly, replication-defective, self-inactivating retroviral constructs were used to establish stable A549 cell lines as previously described. Selection was performed with puromycin (5 μg/mL) for at least 48 hours. Cells were then sorted as a polyclonal population with homogeneous GFP expression and used in subsequent experiments (FACS Aria, IRCAN facility).

### Western blot

Proteins were extracted from cells using Laemmli lysis buffer (12.5 mM Na2HPO4, 15% glycerol, 3% SDS). The protein concentration was measured with the DC Protein Assay (Bio-Rad, Gémenos, France), and total proteins were loaded onto 15% SDS-polyacrylamide gels for electrophoresis and transferred onto polyvinylidene difluoride (PVDF) membranes (Merck Millipore, Guyancourt, France). After 1h of blocking with 3% bovine serum albumin prepared in phosphate-buffered saline (PBS)-0.1% Tween-20 buffer, the blots were incubated overnight at 4°C with primary antibodies: anti-puromycin at 1:10,000 (Millipore Cat# MABE343, RRID: AB_2566826) and anti-α-actinin at 1:10,000 (Millipore, Cat# 05-384, RRID: AB_11212399). After 1h of incubation with a horseradish peroxidase-conjugated secondary antibody (Promega, Charbonnières-les-Bains, France, anti-mouse: Cat# W4021, RRID: AB_430834) at 1:5,000, protein bands were visualized using an enhanced chemiluminescence detection kit (Merck Millipore) and the Syngene Pxi4 imaging system (Ozyme, Saint-Cyr-l’École, France). Immunoblot quantification was performed using Fiji freeware (RRID: SCR_002285) on unsaturated captured images.

### Immunofluorescence, microscopy, and P-body quantification

Cells were grown to subconfluence and fixed at 37°C in 4% paraformaldehyde for 15 min. After fixation, cells were permeabilized with a solution containing 0.3% Triton X-100 for 5 min and incubated with blocking buffer (0.03% Triton X-100, 0.2% gelatin, and 1% BSA) for 20 min. Then, cells were incubated with primary antibodies diluted at 1:200: anti-DDX6 (Novus, distributed by Bio-Techne SAS, Noyal Châtillon sur Seiche (Fr), Cat# NB 200-191, RRID: AB_523228), anti-LSM14A (Santa Cruz Biotechnology Cat# sc-398552, RRID: AB_3099527), anti-LSM14B (Thermo Fisher Scientific Cat# PA5-66371, RRID: AB_2664653), and anti-GR (Santa Cruz Biotechnology, Cat# sc-393232, RRID: AB_2687823) overnight at 4°C in humidified chambers in blocking buffer. Cells were washed and incubated with Alexa Fluor-conjugated secondary antibodies at 1:500 (Thermo Fisher Scientific, Illkirch-Graffenstaden, France, anti-mouse AlexaFluor 594 (Thermo Fisher Scientific Cat# A-21203, RRID: AB_2535789) or AlexaFluor 633 (Thermo Fisher Scientific Cat# A-21050, RRID: AB_2535718), anti-rabbit AlexaFluor 488 (Cat# A-21441, RRID: AB_2535859) or AlexaFluor 594 (Cat# A-21442, RRID: AB_141840) for 1h at room temperature, washed, stained for 5 min with DAPI at 1:10,000 (Thermoscientific, Cat# 62248) and mounted with antifade fluorescence mounting medium (Abcam, Cat# AB104135). Images z-stacks of fluorescence were acquired on an upright epifluorescent microscope (Carl Zeiss) through a 40x/1.3 Plan-Apochromat objective with an ORCA-

Fusion BT digital CMOS Camera (Hamamatsu). Image analysis was conducted on maximum intensity projection images with CellProfiler Image Analysis Software, V2.0 (http://www.cellprofiler.org, RRID: nif-0000-00280). Briefly, all the cells from each image were segmented using the “IdentifyPrimaryObjects” function and the P-bodies were detected using the “IdentifyPrimaryObjects” function and counted per cell using the “MeasureObjectSizeShape” and “RelateObjects” functions.

### Statistical analysis for microscopy analyses

Quantitative data were described and presented graphically as boxplots with medians and standard deviations (immunofluorescence quantification). All statistical analyses were performed using the Prism10 program from GraphPad software (RRID: SCR_002798). The distribution normality was tested with Shapiro’s test. Statistical comparisons were performed using Dunn’s multiple comparison test and considered significant with an alpha level of p < 0.05 (graphically: * for p < 0.05, ** for p < 0.01, *** for p < 0.001, and ns for non-significant).

### Bulk RNA-seq library preparation, sequencing, and analysis

Total RNA was extracted from A549 cells treated with or without 1 μM dexamethasone for 48h (n=4) with TRIzol reagent (Thermo Fisher) and purified on Direct-zol RNA miniprep kit (Zymo Research, Orange, CA). RNA integrity was assessed using an Agilent 2100 Bioanalyzer (RIN ≥8.5). Libraries were prepared using 200 ng of total RNA after ribosomal RNA depletion (Ribo Zero Gold rRNA Removal Kit, Illumina) by Novogene. Fragmentation was carried out using divalent cations under elevated temperature in First Strand Synthesis Reaction Buffer (5X). First-strand cDNA was synthesized using random hexamer primers and M-MuLV Reverse Transcriptase (RNase H-). Second-strand cDNA synthesis was subsequently performed using DNA Polymerase I and RNase H. Remaining overhangs were converted into blunt ends via exonuclease/polymerase activities. After adenylation of the 3’ ends of DNA fragments, adapters with hairpin loop structure were ligated to prepare for hybridization. To select cDNA fragments of preferentially 370-420 bp in length, the library fragments were purified with the AMPure XP system (Beverly, USA). Enzyme (3 μL) was used with size-selected, adaptor-ligated cDNA at 37°C for 15 min, followed by 5 min at 95°C before PCR. PCR was performed using Phusion High-Fidelity DNA polymerase, Universal PCR primers, and Index (X) Primer. PCR products were purified (AMPure XP system), and library quality was assessed using the Agilent 5400 system and quantified by qPCR.

### Sequencing and Data Analysis

Libraries were pooled in equimolar ratios and sequenced on an Illumina NovaSeq 6000 platform using a 150 bp paired-end configuration to a depth of 40-50 million read pairs per sample. Raw reads were quality-checked using FastQC (v0.11.9), adapter sequences were trimmed using Cutadapt (v3.4), and resulting reads were aligned to the human reference genome (GRCh38) using the STAR aligner (v2.7.9a). Gene expression was quantified with feature Counts (Subread package v2.0.3), and differential expression analysis was performed using DESeq2 (v1.32.0) with Phantasus. Genes with fold change ≥1.5 and adjusted p-value <0.01were considered significantly differentially expressed. Pathway enrichment analysis was conducted using GSEA (v4.1.0) with ShinyGO.

### Mass spectrometry analysis

Proteomic analysis was performed on A549 cells cultured with or without 1 μM dexamethasone for 48h (n=4 independent biological replicates per condition). These samples were prepared in parallel with those used for RNA-seq analysis, with duplicate cell cultures processed simultaneously on the same day-one set for transcriptomic analysis and one set for proteomics. This paired experimental design ensured direct comparability between our transcriptomic and proteomic datasets. Total proteins were extracted using Laemmli buffer and processed for LC MS/MS analysis at the IGBMC proteomic facility. For protein digestion, 30 μg of protein per sample was precipitated with TCA overnight at 4°C, followed by two cold acetone washes. Dried pellets were dissolved in 1 M urea in 0.1 mM TrisHCl (pH 8.5) for subsequent reduction with 10 mM DTT (30 min, 56°C) and alkylation with 20 mM iodoacetamide (30 min, 25°C). Trypsin digestion was performed with two sequential enzyme additions (initial and at 2h) for overnight digestion at 37°C. Samples were acidified with 0.2% TFA before MS analysis. NanoLC-MS/MS was performed using an Ultimate 3000 nano-RSLC system coupled to an Exploris 480 quadrupole-orbitrap via a nano-electrospray source with FAIMS pro interface (Thermo Fisher Scientific). Tryptic peptides (1 μl) were preconcentrated on a C18 PepMap100 trap column (300 μm × 1 mm) for 1 min at 15 μL/min with 2% ACN and 0.1% FA in water. Peptide separation was achieved on an analytical column (C18 PepMap, 75 μm ID × 15 cm) using a 40-min gradient from 8% to 25% buffer B (buffer A: 0.1% FA in water; buffer B: 0.1% FA in 80% ACN) at 450 nl/min and 45°C. The gradient was followed by a regeneration step at 90% B and re-equilibration to 8% B, with a total chromatography time of 120 min. The mass spectrometer operated in positive ionization mode using Data-Dependent Acquisition with two cycles of FAIMS compensation voltages (−45V and −55V for 1.2 and 0.8 sec, respectively). Each FAIMS-DDA cycle consisted of a survey scan (350-1200 m/z, 60,000 FWHM) followed by MS^2^ spectra acquisition (HCD; 30% normalized energy; 2 m/z window; 22,500 FWHM). Normalized AGC values were set at 300% for MS1 and 100% for MS2, with 50 ms maximum injection time for both scan modes. Single-charged and unassigned states were excluded, with a 40 s exclusion duration (± 10 ppm mass width).

### MS data processing

Proteins were identified using Proteome Discoverer 2.5 software (Thermo Scientific) against the Human proteome database (SwissProt, reviewed, March 2024 release). Precursor and fragment mass tolerances were set at 7 ppm and 0.05 Da, respectively, allowing up to two missed cleavages. Oxidation (M) was set as a variable modification and carbamidomethylation (C) as a fixed modification, with peptide filtering at 1% FDR. Protein quantification required a minimum of one unique peptide based on XIC values. The mass spectrometry proteomics data have been deposited to the ProteomeXchange Consortium via the PRIDE37 582 partner repository with the dataset identifier PXD063431.

Subsequent bioinformatic analysis was performed in R using the ‘wrProteo’ (https://CRAN.R-project.org/package=wrProteo; doi:10.32614/CRAN.package.wrProteo) package, including data filtering (removing proteins with ≤3 PSM or detected in <70% of samples), variance stabilization, and missing value imputation using the k-nearest neighbor method (wrProteo). Differential protein expression between dexamethasone-treated and untreated A549 cells was determined using linear models with empirical Bayes statistics. Proteins with absolute log_2_ fold change ≥1.2 and adjusted p-value ≤0.05 (Benjamini-Hochberg correction) were considered significantly differentially expressed. For correlation analysis, genes with corresponding Ensembl identifiers were filtered. From these genes, we generated lists of fold change values for transcripts and proteins following dexamethasone treatment, which were plotted on the same graph to assess the correlation between mRNA and protein level changes 48 hours post-treatment. GC-content was extracted from Ensembl identifiers for cDNA, 5’UTR, coding region (CDS), and 3’UTR.

## Data availability

The authors declare that all data supporting the findings of this study are available within the article, and its supplementary data file. Whole transcriptomic data was similarly deposited under accession number GSE295999 and the proteomic data accession number Pride (PXD063431).

## Acknowledgements

Acknowledgments Authors would like to thank Dr. Julien Cherfils and Ludovic Cervera (Cytomed facility), and Dr. Soline Estrach and Dr. Aaron Mendez-Bermudez (PICMI), Dr. Christian Baudoin (GENOMED), Dr. Luc Negroni for technical help, and also some collaborators such as, Dr. Dominique Weil and Dr. Michel Kress, Pr. Franck Delaunay for plasmid kind gifts and Dr. David Gilot for the Mel501 cells as well as Dr. Hubstenberger’s team member for the scientific discussions. We also thank all members from the team from the Centre Scientifique de Monaco who helped us generating the CRISPR cell lines. This work has been supported by grants from the French Government (Agence Nationale de Recherche, ANR) through the “Investments for the Future” LABEX SIGNALIFE (ANR-11-LABX 0028-01, IDEX UCAJedi ANR-15-IDEX-01, under the France 2030 programme, reference ANR-23-IAHU-0007); “Fondation ARC pour la Recherche contre le Cancer” (Canc’air GENEx posomics and PJA-20191209562), Canceropole PACA, Institut National du Cancer PL-Bio (INCa_18414), INSERM cancer; Plan Cancer 2014-2019 (18CN045). The authors acknowledge PICMI and Cytomed, the IRCAN’s Imaging facility core part of the “Microscopie Imagerie Cytométrie Azur” (MICA) GIS IBiSA labeled platform. PICMI and Cytomed were supported by the Association pour la Recherche sur le Cancer (ARC), by the national research infrastructures in biology and health (GIS IBiSA) as part of MICA and by the “Conseil Départemental 06 et Région Sud”. PICMI and Cytomed were supported by FEDER, Contrat Plan Etat Région (CPER), Ministère de l’Enseignement Supérieur et de la Recherche, Région Sud, Conseil Départemental 06, ITMO Cancer Aviesan (Plan Cancer), Cancéropole PACA, INCa, CNRS and Inserm. PCBIS thanks the Strasbourg Drug Discovery and Development Institute (IMS), as part of the Interdisciplinary Thematic Institute (ITI) 2021–2028 program of the University of Strasbourg, CNRS, and Inserm, supported by IdEx Unistra (ANR-10-IDEX-0002) and 23 618 by the SFRI-STRAT’US project (ANR-20-SFRI-0012) under the framework of the French Investments for the Future Program. IGBMC proteomic facility was supported by IGBMC, CNRS, Inserm and Université de Strasbourg.

## Contributions

Conceptualization: V.J.N., M.D., K.J. and P.B.; Methodology: V.J.N., K.J., M.I., O.V-C., J.D. and P.B.; Software: W.R., F.B., S.A.; Validation: V.J.N. and P.B.; Formal analysis: V.J.N., A.B., W.R., A.K., B.D., P.V., C.DG. and P.B.; Investigation: V.J.N., M.D., K.J., O.V-C., T.R., J.D., C.G., and A.K.; Resources: A.H., P.H. and P.B.; Data curation: V.J.N. and P.B.; Writing-original draft preparation: V.J.N., P.H. and P.B.; Writing-review and editing: V.J.N., M.D., B.M., P.H., A.H. and P.B.; Visualization: V.J.N. and P.B.; Supervision: P.B.; Project administration; V.J.N. and P.B.; Funding acquisition: V.J.N., A.H., P.H., and P.B. All authors have read and agreed to the published version of the manuscript.

