## Supplementary Figures for "Glucocorticoids Modulate mRNA Translation Fate Through P-Body Dynamics"

#### Supplementary Figure legends

**Supplementary Figure 1: Screening and structural analysis of glucocorticoids as P-body regulators. (A)** Table summarizing drug screening results, displaying P-body fold change, cell number fold change, and P-body to cell number ratio every 4 hours for 72h. **(B)** UMAP similarity analysis filtered through pharmacophore classification. Path fingerprints were extracted from chemical structures of the 98 steroids in the Prestwick library. These fingerprints, which capture bonding paths between atoms, were used to identify common substructures and recurring molecular patterns. The UMAP dimension reduction algorithm was applied to these fingerprints to generate a 3D representation where compounds are positioned according to their structural similarity. The resulting similarity graph, combined with activity data, provides a comprehensive mapping of the steroid chemical space. **(C)** Molecular substructure of dexamethasone and prednisolone. **(D)** Confusion matrix displaying true positives (TPs), false positives (FPs), true negatives (TNs), and false negatives (FNs) for each class. Precision values quantify the proportion of correct positive predictions among all positive predictions made.

**Supplementary Figure 2: Identification of glucocorticoids as positive regulators of P-bodies. (A-C)** Representative immunofluorescence images of DDX6 (P-body marker, green) in **(A)** A549, **(B)** HeLa and **(C)** Mel501 cells treated for 48h with 1 μM of either Dexa or Pred. Cells were additionally treated with sodium arsenite ($NaAsO_2$, 0.5 mM) for 30 min to induce P-body formation. Scale bar = 10 μm. **(D-E-F)** Quantification of **(D)** P-body number per cell, **(E)** P-body mean area per cell and **(F)** P-body total area (in fold change) under experimental conditions shown in panel A, from three independent biological experiments (number of cells > 100/condition).

**Supplementary Figure 3: Differential effects of glucocorticoids on P-bodies and stress granules. (A)** Representative immunofluorescence images of LSM14A (P-body marker, red) in GFP-DDX6 A549 cells treated with 0.1 and 1 μM of dexamethasone (Dexa) or prednisolone (Pred). Scale bar = 10 μm. **(B)** Quantification of P-body number per cell under experimental conditions shown in panel A, from three independent biological experiments (number of cells > 100/condition). **(C-E)** Representative immunofluorescence images of DDX6 (P-body marker) in **(C)** A549, **(D)** HeLa and **(E)** Mel501 cells treated with 0.1 μM of dexamethasone or prednisolone. Scale bar = 10 μm.

**Supplementary Figure 4: Glucocorticoids regulate specifically P-bodies. (A-C-E)** Representative immunofluorescence images of DDX6 (green) and glucocorticoid receptor (GR, red) in **(A)** A549, **(C)** HeLa and **(E)** Mel501 cells treated for 48h with 0.1 μM of dexamethasone (Dexa), or 0.1 μM of prednisolone (Pred), and/or 1 μM of RU486. Scale bar = 10 μm. **(B-D-F)** Quantification of P-body number per cell under experimental conditions shown in panels A, C and E respectively, from three independent biological experiments (number of cells > 100/condition).

**Supplementary Figure 5: Glucocorticoid receptor mediates P-body formation in multiple cell lines. (A-C-E)** Immunoblot analysis of GR, phosphorylated GR at serine 211 (pGR S211), FKBP5, and GAPDH proteins from **(A)** A549, **(C)** HeLa and **(E)** Mel501 cells treated for 48h with 0.1 µM of Dexa, or Pred and/or 1 µM of RU486. **(B-D-F)** Quantification of the immunoblot from panels A, B and C respectively, from three independent experiments.

**Supplementary Figure 6: Glucocorticoid receptor mediates P-body formation in KO GR cells. (A)** Representative immunofluorescence images of DDX6 (green) and GR (red) in control A549 cells (A549 CTL) and two CRISPR-mediated GR knockout A549 clones (A549 KO GR#1 and #2) treated for 48h with 0.1 µM of Dexa. Scale bar = 10 µm. **(B)** Quantification of P-body number under experimental conditions shown in panel A, from three independent biological experiments (number of cells > 100/condition). **(C)** Immunoblot analysis of GR, pGR S211, FKBP5 and GAPDH in control A549 cells (CTL) and A549 KO GR#1 and #2 cells treated for 48h with 0.1 µM of Dexa.

**Supplementary Figure 7: Glucocorticoid Receptor** a **Isoform is important for P-body regulation. (A)** Representative immunofluorescence images of DDX6 (red) and GFP constructs (green) in A549 GFP or KO GR#1 cells complemented with GFP or GFP-GRa and treated for 48h with 0.1 µM of dexamethasone (Dexa). Scale bar = 10 µm. **(B)** Quantification of P-body number under experimental conditions shown in panel A, from three independent biological experiments (number of cells > 100/condition). **(C)** Immunoblot analysis of GR with two bands: one for the endogenous (endo) and one of the GR fused to GFP, phosphorylated GR at serine 211 (pGR S211) with two bands: one for the endogenous (endo) and one of the pGR fused to GFP, FKBP5, and GAPDH proteins from cells expressing different GFP constructs in A549 or KO GR#1 cells complemented with GFP or GFP-GRa and treated with Dexa from 0.1 nM to 1 µM. **(D)** Representative immunofluorescence images of G3BP1 (red) in A549 GFP and the two different KO GR clones complemented with GFP-GRa cells treated with Dexa from 0.1 nM to 1 µM. Scale bar = 10 µm.

**Supplementary Figure 8: Characterization of GR constructs for P-body regulation. (A)** Schematic representation of the different GR constructions (created with https://BioRender.com/zwtur4m). **(B)** Immunoblot analysis of GR, phosphorylated GR at serine 211 (pGR S211), FKBP5, and GAPDH proteins from A549 CTL and A549 KO GR#1 and #2 cells complemented with the different GR constructions, treated for 48h with 0.1 µM of Dexa and/or 1 µM of RU486. **(C)** Representative immunofluorescence images of DDX6 (red) and the GFP constructions (green) in A549 KO GR#1 complemented with GFP-GRb or GFP-GRΔEx9 cells treated for 48h with 0.1 µM of Dexa. Scale bar = 10 µm. **(D)** Quantification of P-body number per cell under experimental conditions shown in panel C, from three independent biological experiments (number of cells > 100/condition).

**Supplementary Figure 9: GR isoform specificity in P-body regulation. (A)** Representative immunofluorescence images of DDX6 (red) and the GFP constructions (green) in A549 KO GR#2 complemented with GFP, GFP-GRa, GFP-GRb or GFP-GRΔEx9 cells treated for 48h with 0.1 µM of Dexa. Scale bar = 10 µm. **(B)** Quantification of P-body number per cell under experimental conditions shown in panel A, from three independent biological experiments (number of cells > 100/condition). **(C)** Representative immunofluorescence images of LSM14A (red) in A549 KO GR#2 complemented with GFP-GRa cells treated for 48h with 0.1 µM of Dexa. Scale bar = 10 µm. **(D)** Quantification of P-body number per cell under experimental conditions shown in panel C, from three independent biological experiments (number of cells > 100/condition).

**Supplementary Figure 10: GR activation enriches P-body mRNA targets but alters their translation. (A)** Volcano plot showing differential mRNA expression in dexamethasone-treated versus untreated A549 cells. Purple dots represent transcripts previously identified as P-body-enriched. Statistical significance (-log10 p-value) is plotted against magnitude of change (log2 fold change). **(B)** Volcano plot depicting differential protein expression in dexamethasone-treated versus untreated A549 cells. Purple dots represent proteins encoded by mRNAs previously identified as P-body-enriched. Statistical significance (-log10 p-value) is plotted against magnitude of change (log2 fold change). **(C)** Gene Set Enrichment Analysis (GSEA) previously established glucocorticoid receptor-induced gene signatures within differentially expressed mRNAs following dexamethasone treatment. Nominal p-values were calculated using GSEA software. **(D)** Gene Set Enrichment Analysis (GSEA) previously established glucocorticoid receptor-induced gene signatures within differentially expressed proteins following dexamethasone treatment. Nominal p-values were calculated using GSEA software. **(E)** Gene Set Enrichment Analysis (GSEA) of P-body-associated transcripts within differentially expressed mRNAs following dexamethasone treatment. Nominal p-values were calculated using GSEA software. **(F)** Gene Set Enrichment Analysis (GSEA) of P-body-associated transcripts within differentially expressed proteins following dexamethasone treatment. Nominal p-values were calculated using GSEA software. **(G)** Gene Set Enrichment Analysis (GSEA) of ENCODE transcription factor signature in differentially expressed genes following dexamethasone treatment. Nominal p-values were calculated using GSEA software. **(H)** Gene Set Enrichment Analysis (GSEA) of ENCODE transcription factor signature in differentially expressed proteins following dexamethasone treatment. Nominal p-values were calculated using GSEA software. **(I)** Correlation plot of protein abundance versus corresponding mRNA levels. Purple dots represent proteins encoded by mRNAs previously identified as P-body-enriched, demonstrating disrupted correlation between transcriptome and proteome for this subset. **(J)** Gene Set Enrichment Analysis of transcripts exhibiting decreased translational efficiency (reduced protein ratio) following dexamethasone treatment. Functional enrichment analysis was performed using ShinyGO software.

**Supplementary Figure 11: Glucocorticoid receptor promotes P-body formation through LSM14B regulation.** **(A)** LSM14A relative RNA and protein expression (in fold change) from A549 cells treated with 1 µM of dexa for 48 hours. **(B)** LSM14B RNA and protein relative expression (in fold change) from A549 cells treated with 1 µM of dexa for 48 hours. **(C)** Representative immunofluorescence images of LSM14A (yellow) and LSM14B (red) in A549 cells treated with Dexa at 0.1 µM for 48 hours. Scale bar = 10 µm. **(D)** Immunoblot analysis of GR, phosphorylated GR at serine 211 (pGR S211), FKBP5, LSM14B, and GAPDH proteins from A549 cells treated with Dexa at 0.1 µM for 48 hours. **(E)** Representative immunofluorescence images of LSM14B (green) and LSM14A (red) in A549 cells transfected with siCTL or two different siRNAs against LSM14B. Scale bar = 10 µm. **(F)** Immunoblot analysis of LSM14B, LSM14A, GR, and GAPDH proteins from A549 cells transfected with siCTL or two different siRNAs against LSM14B. **(G)** Immunoblot analysis of LSM14B with two bands: one for the endogenous (endo) and one for LSM14B fused to GFP, GR, phosphorylated GR at serine 211 (pGR S211), FKBP5, and GAPDH proteins from A549 cells expressing GFP or GFP-LSM14B treated with 0.1 µM of Dexa for 48 hours. **(H)** Relative protein expression (in fold change) of LSM14B and c-JUN from A549 cells treated with 10µg/ml of cycloheximide at different time point.

**Supplementary Figure 12: LSM14B is regulated by glucocorticoid receptor across cell lines.** **(A)** Representative immunofluorescence images of LSM14A (yellow) and LSM14B (red) in A549, HeLa or Mel501 cells treated with 0.1 µM of Dexa for 48 hours. Scale bar = 10 µm. **(B)** Quantification of P-body number per cell under experimental conditions shown in panel A, from three independent biological experiments (number of cells > 100/condition). **(C)** Immunoblot analysis of GR, phosphorylated GR at serine 211 (pGR S211), FKBP5, LSM14B, and GAPDH proteins from A549 cells treated with 0.1 µM of Dexa and/or 1 µM of RU486 or relacorilant (Rela: another GR inhibitor) for 48 hours. **(D)** Quantification of the immunoblot from panel C, from three independent experiments. **(E)** Immunoblot analysis of GR, phosphorylated GR at serine 211 (pGR S211), FKBP5, LSM14B, and GAPDH proteins from A549 CTL, KO GR#1 and KO GR#1+GRa cells treated with 0.1 µM of Dexa and/or 1 µM of RU486 48 hours. **(F)** Quantification of the immunoblot from panel E, from three independent experiments.

**Supplementary Figure 13: Glucocorticoid-induced P-body formation is reversible upon glucocorticoid withdrawal.** **(A)** Representative immunofluorescence images of LSM14A (red) and LSM14B (green) in A549 cells treated with 0.1 µM of Dexa (no withdrawal), after 48h of treatment medium was changed to normal medium for 24h (withdrawal). Scale bar = 10 µm. **(B)** Representative immunofluorescence images of DDX6 (yellow) in A549 KO GR#1 and

KO GR#1 + GRa cells treated with 0.1 µM of Dexa (no washout), after 48h of treatment medium was changed to normal medium for 24h (washout). Scale bar = 10 µm. **(C)** Quantification of P-body number under experimental conditions shown in panel B, from three independent biological experiments (number of cells > 100/condition). **(D)** Representative immunofluorescence images of DDX6 (yellow) in A549 KO GR#2 and KO GR#2 + GRa cells treated with 0.1 µM of Dexa (no washout), after 48h of treatment medium was changed to normal medium for 24h (washout). Scale bar = 10 µm. **(E)** Quantification of P-body number under experimental conditions shown in panel D, from three independent biological experiments (number of cells > 100/condition).

**A**

| Name | Family | P-Body fold change 0 h | 24 h | 48 h | 72 h | Confluency fold change 0 h | 24 h | 48 h | 72 h | P-Body/Confluency fold change 0 h | 24 h | 48 h | 72 h |
|---|---|---|---|---|---|---|---|---|---|---|---|---|---|
| Fluticasone propionate 1µM | **Glucocorticoid** | 103% | 167% | 250% | 213% | 111% | 116% | 96% | 75% | 96% | 137% | 276% | 331% |
| Fluticasone propionate 10µM | **Glucocorticoid** | 110% | 183% | 269% | 216% | 119% | 126% | 102% | 80% | 95% | 137% | 278% | 316% |
| Clocortolone pivalate 1µM | **Glucocorticoid** | 104% | 179% | 273% | 219% | 109% | 119% | 107% | 90% | 98% | 142% | 270% | 282% |
| Prednisolone 10µM | **Glucocorticoid** | 104% | 166% | 238% | 185% | 110% | 122% | 92% | 67% | 93% | 133% | 261% | 279% |
| Fludrocortisone acetate 10µM | **Glucocorticoid** | 130% | 168% | 258% | 211% | 120% | 138% | 107% | 79% | 107% | 119% | 244% | 270% |
| Dexamethasone 1µM | **Glucocorticoid** | 120% | 156% | 230% | 184% | 110% | 125% | 95% | 70% | 108% | 121% | 242% | 266% |
| Prednisolone 1µM | **Glucocorticoid** | 100% | 162% | 244% | 184% | 112% | 124% | 98% | 71% | 88% | 127% | 252% | 261% |
| Dexamethasone 10µM | **Glucocorticoid** | 118% | 142% | 223% | 175% | 113% | 125% | 91% | 68% | 102% | 111% | 245% | 259% |
| Triamcinolone 10µM | **Glucocorticoid** | 121% | 157% | 238% | 189% | 108% | 123% | 97% | 75% | 111% | 124% | 246% | 254% |
| Prednicarbate 1µM | **Glucocorticoid** | 115% | 186% | 280% | 223% | 127% | 136% | 126% | 104% | 93% | 129% | 234% | 249% |
| Clocortolone pivalate 10µM | **Glucocorticoid** | 113% | 173% | 207% | 142% | 107% | 117% | 91% | 67% | 108% | 140% | 239% | 246% |
| Rimexolone 10µM | **Glucocorticoid** | 116% | 193% | 278% | 206% | 118% | 129% | 118% | 100% | 101% | 142% | 250% | 241% |
| Prednicarbate 10µM | **Glucocorticoid** | 120% | 149% | 239% | 203% | 120% | 133% | 122% | 98% | 102% | 106% | 207% | 241% |
| Cortisol acetate 1µM | **Glucocorticoid** | 104% | 175% | 251% | 168% | 102% | 110% | 91% | 68% | 104% | 156% | 271% | 237% |
| Triamcinolone 1µM | **Glucocorticoid** | 132% | 171% | 261% | 209% | 119% | 137% | 113% | 90% | 109% | 122% | 232% | 236% |
| Rimexolone 1µM | **Glucocorticoid** | 104% | 181% | 273% | 221% | 115% | 126% | 124% | 110% | 93% | 136% | 233% | 234% |
| Melengestrol acetate 1µM | **Glucocorticoid** | 106% | 168% | 217% | 180% | 106% | 111% | 102% | 90% | 102% | 144% | 224% | 233% |
| Loteprednol etabonate 1µM | **Glucocorticoid** | 101% | 156% | 208% | 155% | 101% | 110% | 88% | 69% | 102% | 143% | 241% | 223% |
| Tiratricol 1µM | Iodophenylalanines | 123% | 198% | 320% | 366% | 112% | 124% | 145% | 167% | 108% | 156% | 222% | 222% |
| Beclomethasone dipropionate 1µM | **Glucocorticoid** | 99% | 143% | 212% | 144% | 105% | 111% | 84% | 67% | 99% | 132% | 253% | 215% |
| Fludrocortisone acetate 1µM | **Glucocorticoid** | 124% | 177% | 258% | 216% | 114% | 132% | 118% | 102% | 107% | 131% | 221% | 214% |
| Desonide 1µM | **Glucocorticoid** | 102% | 136% | 196% | 139% | 105% | 113% | 90% | 70% | 99% | 117% | 220% | 212% |
| Tiratricol 10µM | Iodophenylalanines | 117% | 181% | 289% | 323% | 110% | 122% | 137% | 155% | 105% | 145% | 213% | 211% |
| Liothyronine 1µM | Iodophenylalanines | 94% | 194% | 279% | 259% | 106% | 106% | 115% | 123% | 95% | 188% | 243% | 211% |
| Fluorometholone 1µM | **Glucocorticoid** | 91% | 138% | 205% | 144% | 103% | 110% | 84% | 69% | 93% | 129% | 246% | 210% |
| Cortisol acetate 10µM | **Glucocorticoid** | 103% | 167% | 231% | 148% | 114% | 116% | 95% | 69% | 91% | 142% | 240% | 206% |
| (L) Thyroxine 1µM | Iodophenylalanines | 92% | 159% | 220% | 229% | 91% | 88% | 98% | 107% | 101% | 180% | 222% | 205% |
| Liothyronine 10µM | Iodophenylalanines | 79% | 169% | 260% | 237% | 96% | 98% | 107% | 117% | 87% | 178% | 243% | 204% |
| Ciclesonide 1µM | **Glucocorticoid** | 98% | 143% | 198% | 132% | 100% | 110% | 86% | 66% | 103% | 134% | 230% | 201% |
| Dichlorisone acetate 1µM | **Glucocorticoid** | 97% | 130% | 180% | 128% | 102% | 109% | 86% | 70% | 97% | 116% | 211% | 195% |
| Desonide 10µM | **Glucocorticoid** | 106% | 114% | 169% | 123% | 99% | 111% | 87% | 69% | 109% | 100% | 196% | 190% |
| Melengestrol acetate 10µM | **Glucocorticoid** | 108% | 156% | 217% | 166% | 113% | 120% | 114% | 104% | 97% | 124% | 201% | 186% |
| (L) Thyroxine 10µM | Iodophenylalanines | 108% | 132% | 195% | 197% | 85% | 89% | 94% | 102% | 128% | 149% | 203% | 185% |
| Corticosterone 1µM | **Glucocorticoid** | 104% | 141% | 198% | 161% | 98% | 100% | 88% | 88% | 106% | 142% | 219% | 174% |
| Triflusal 10µM | Single | 109% | 133% | 190% | 177% | 92% | 99% | 102% | 104% | 121% | 131% | 186% | 170% |
| Dichlorisone acetate 10µM | **Glucocorticoid** | 124% | 101% | 146% | 103% | 126% | 112% | 87% | 68% | 100% | 88% | 169% | 162% |
| Nelarabine 10µM | Adenosine-like | 110% | 133% | 155% | 210% | 111% | 113% | 140% | 159% | 101% | 111% | 117% | 154% |
| Monobenzone 10µM | Single | 100% | 118% | 144% | 160% | 104% | 99% | 112% | 103% | 99% | 117% | 126% | 149% |

**B**

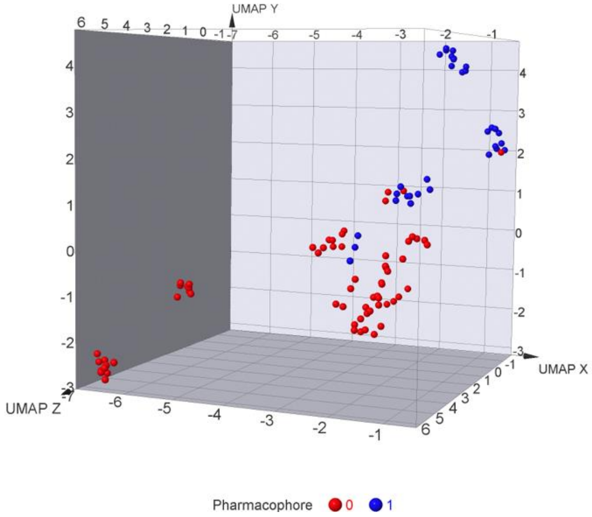


**D**

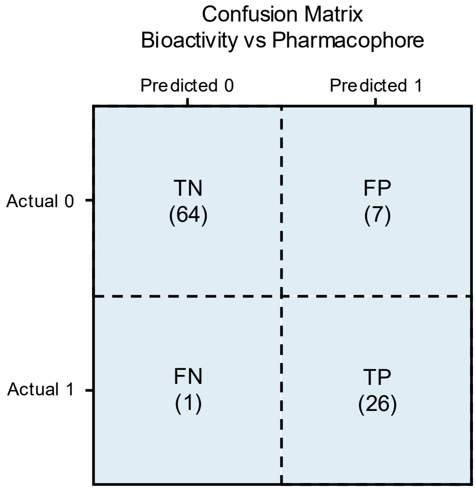


**C**

**Dexamethasone** **Prednisolone**

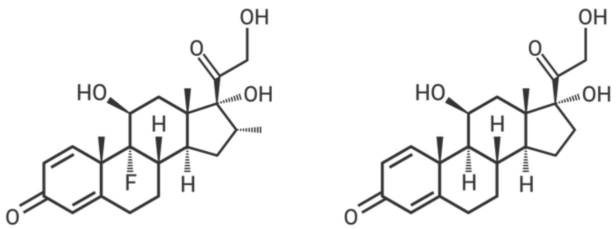

### Supp Figure 1

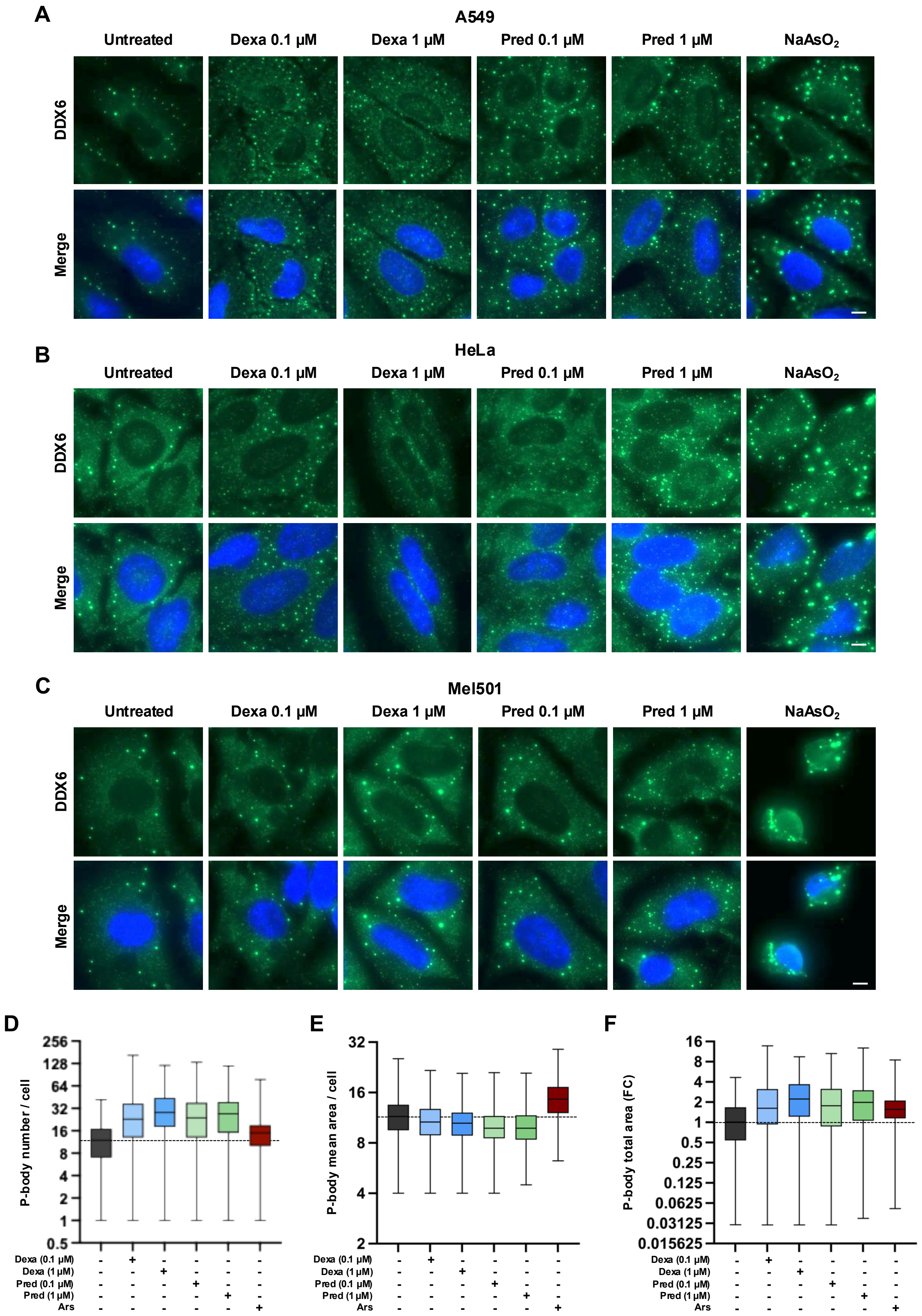


### Supp Figure 2

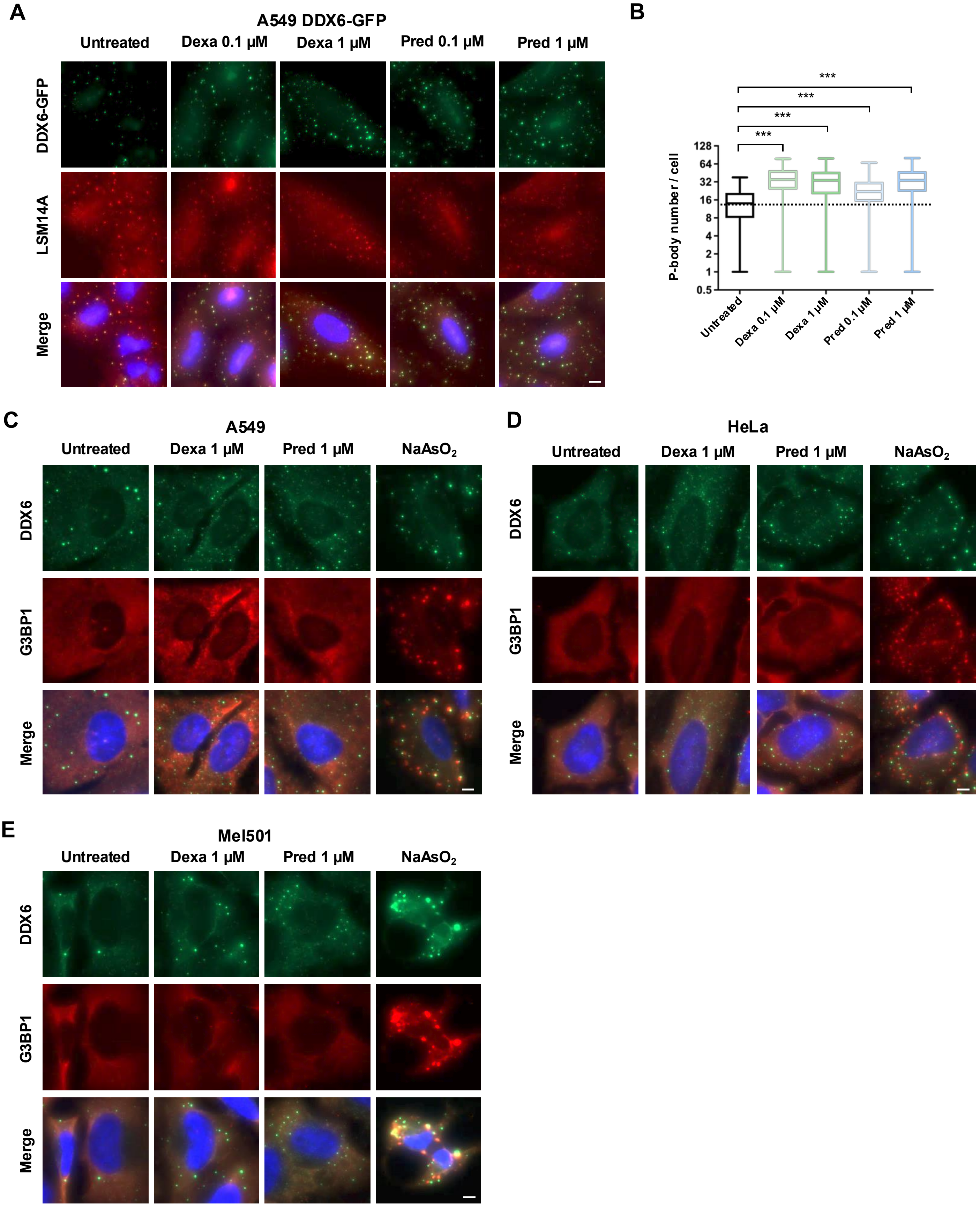


### Supp Figure 3

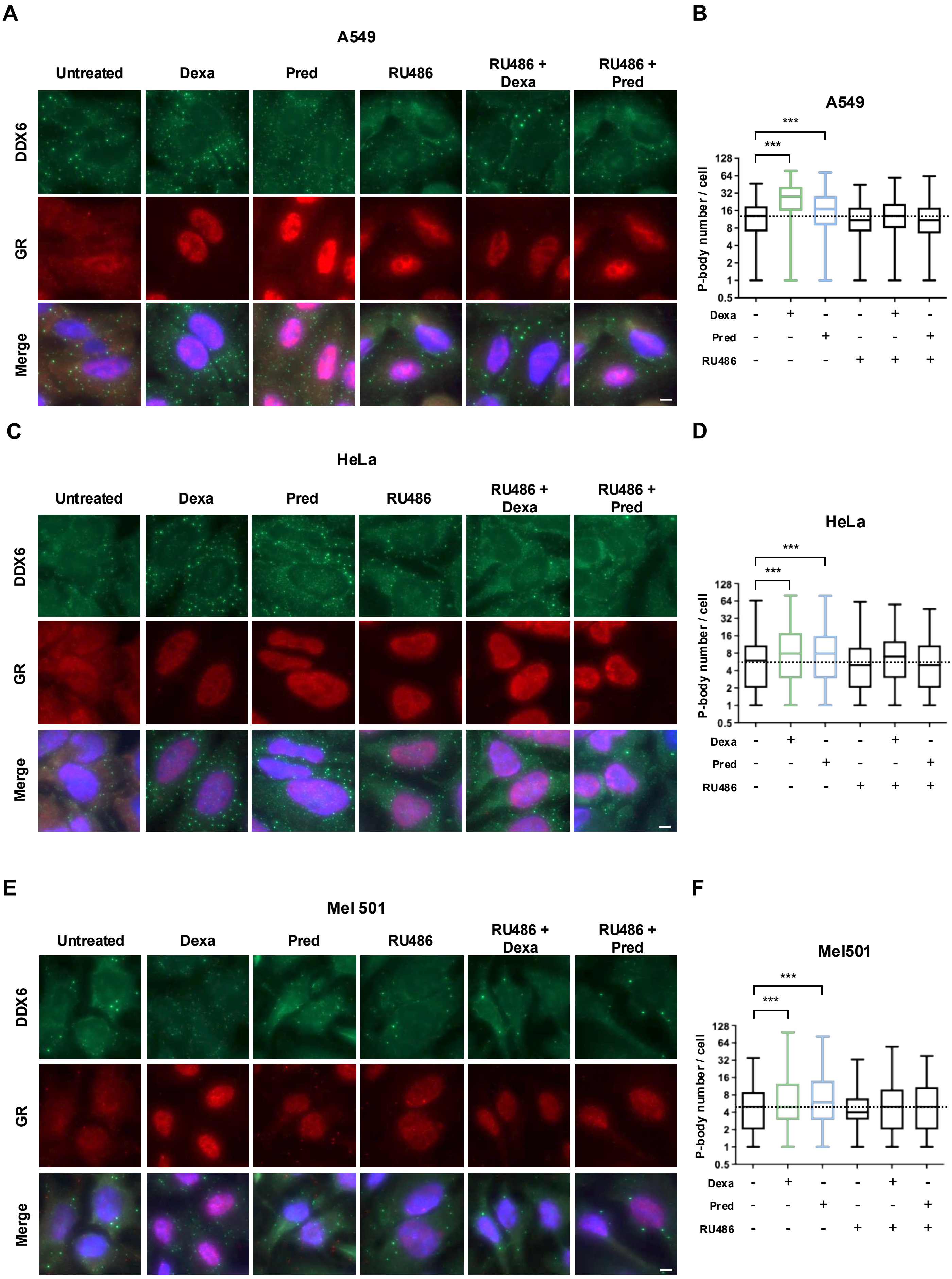


### Supp Figure 4

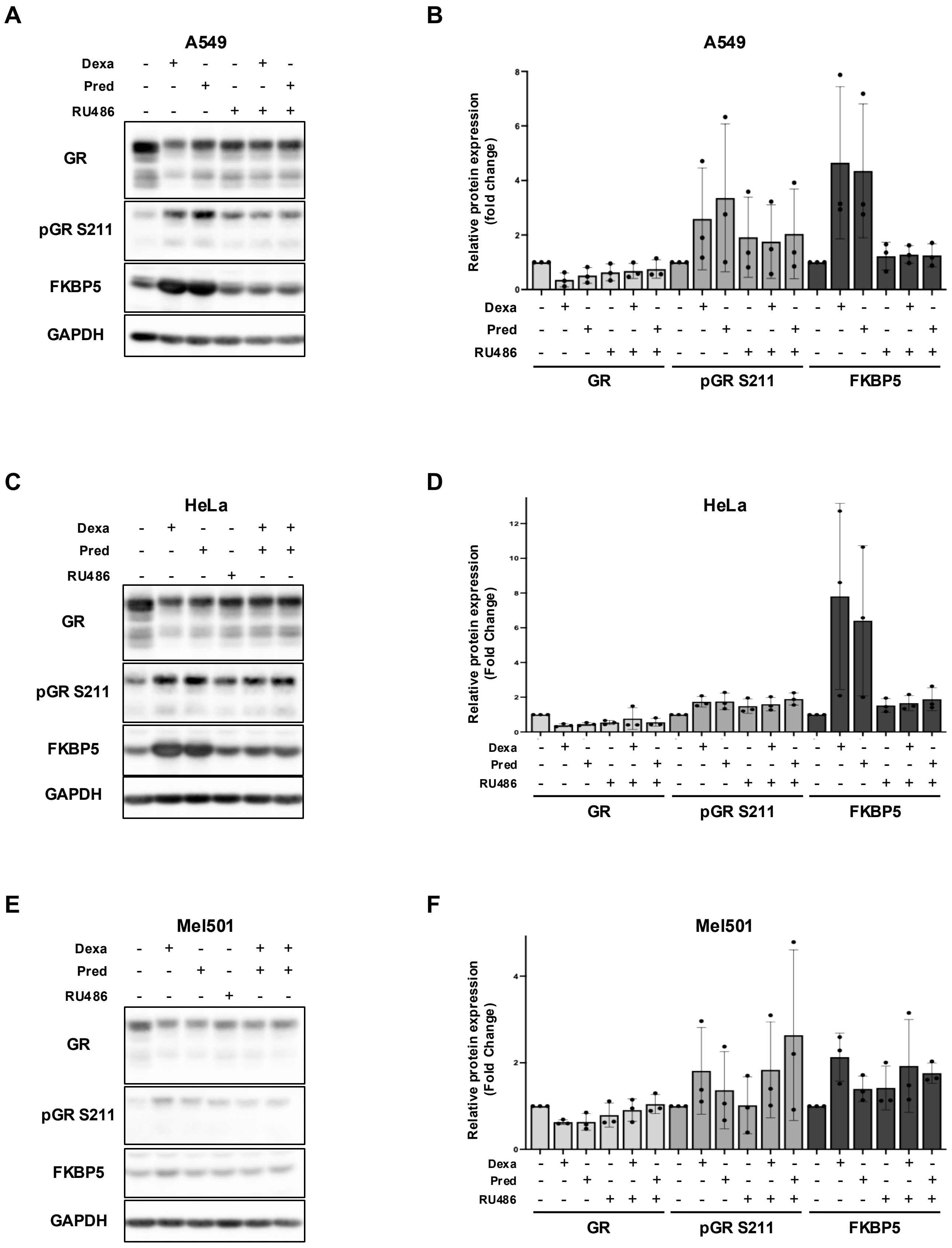


**Supp Figure 5**

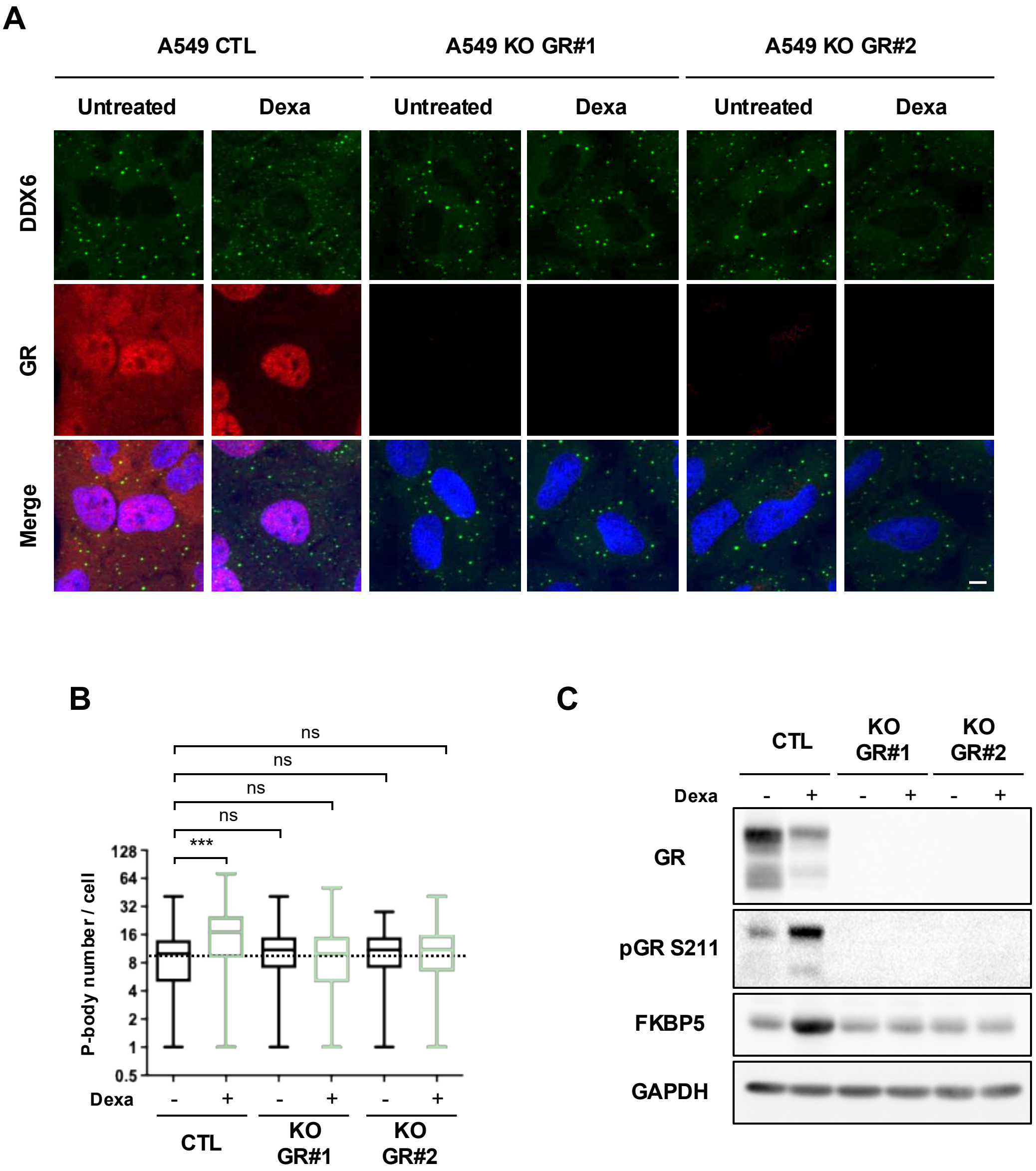


**Supp Figure 6**

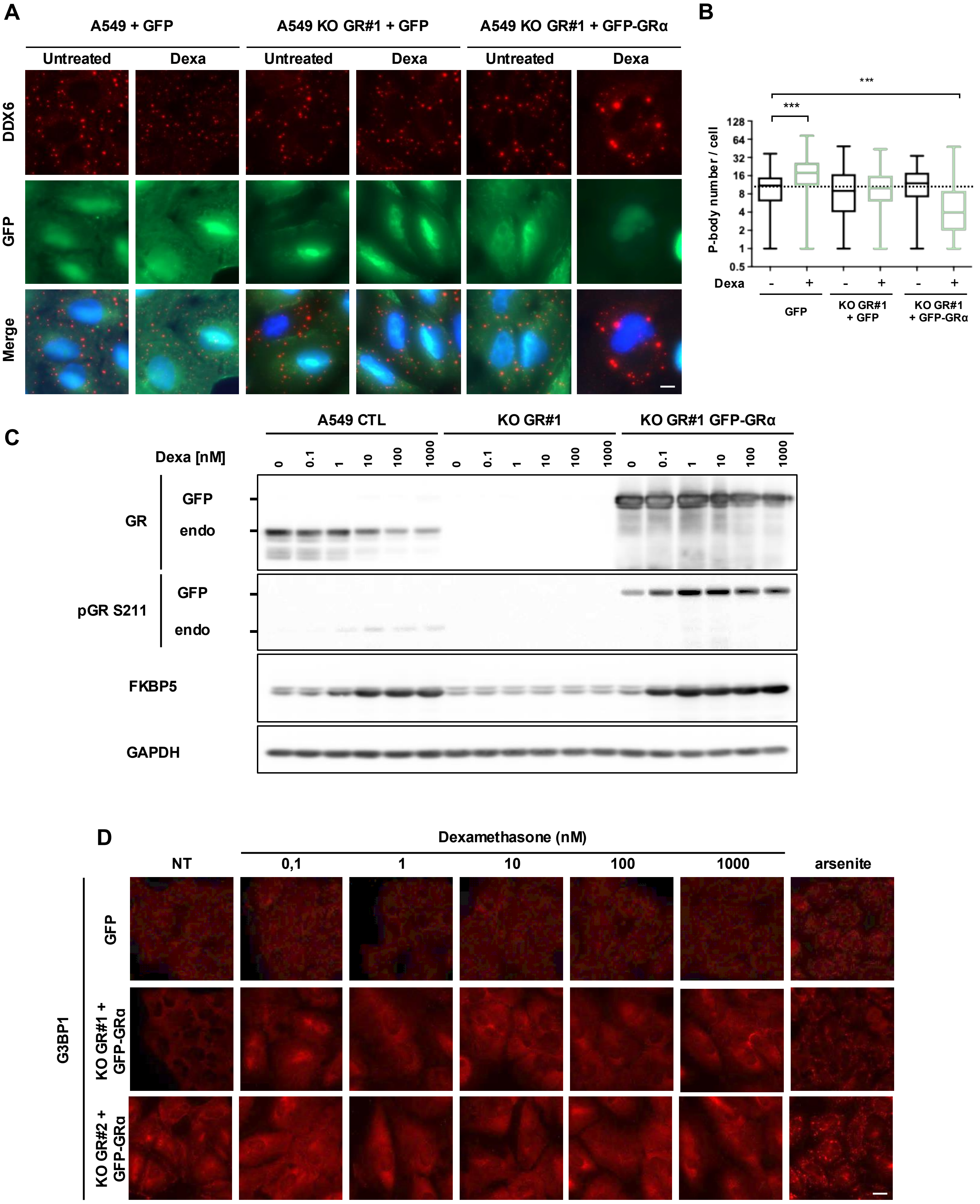


### Supp Figure 7

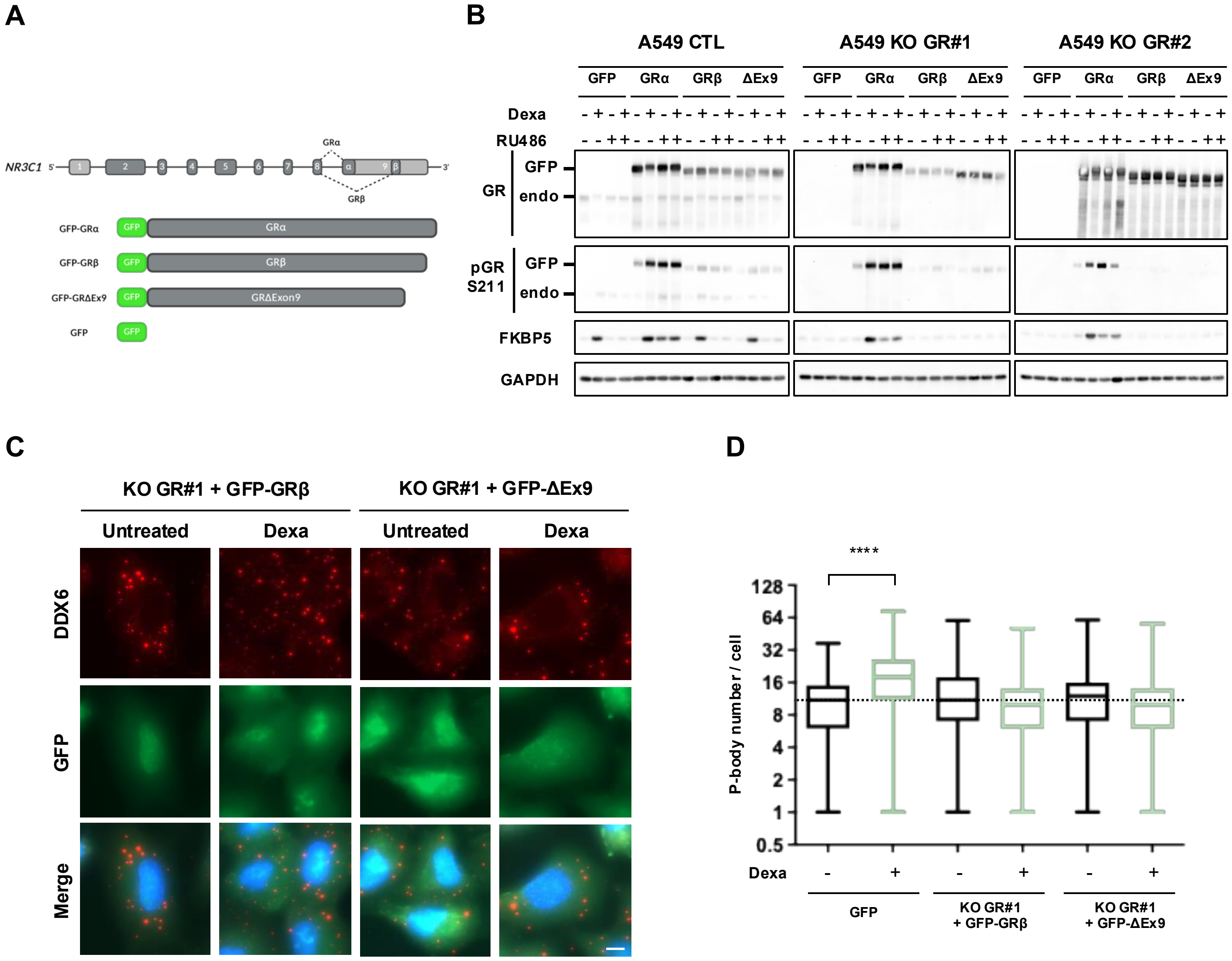


### Supp Figure 8

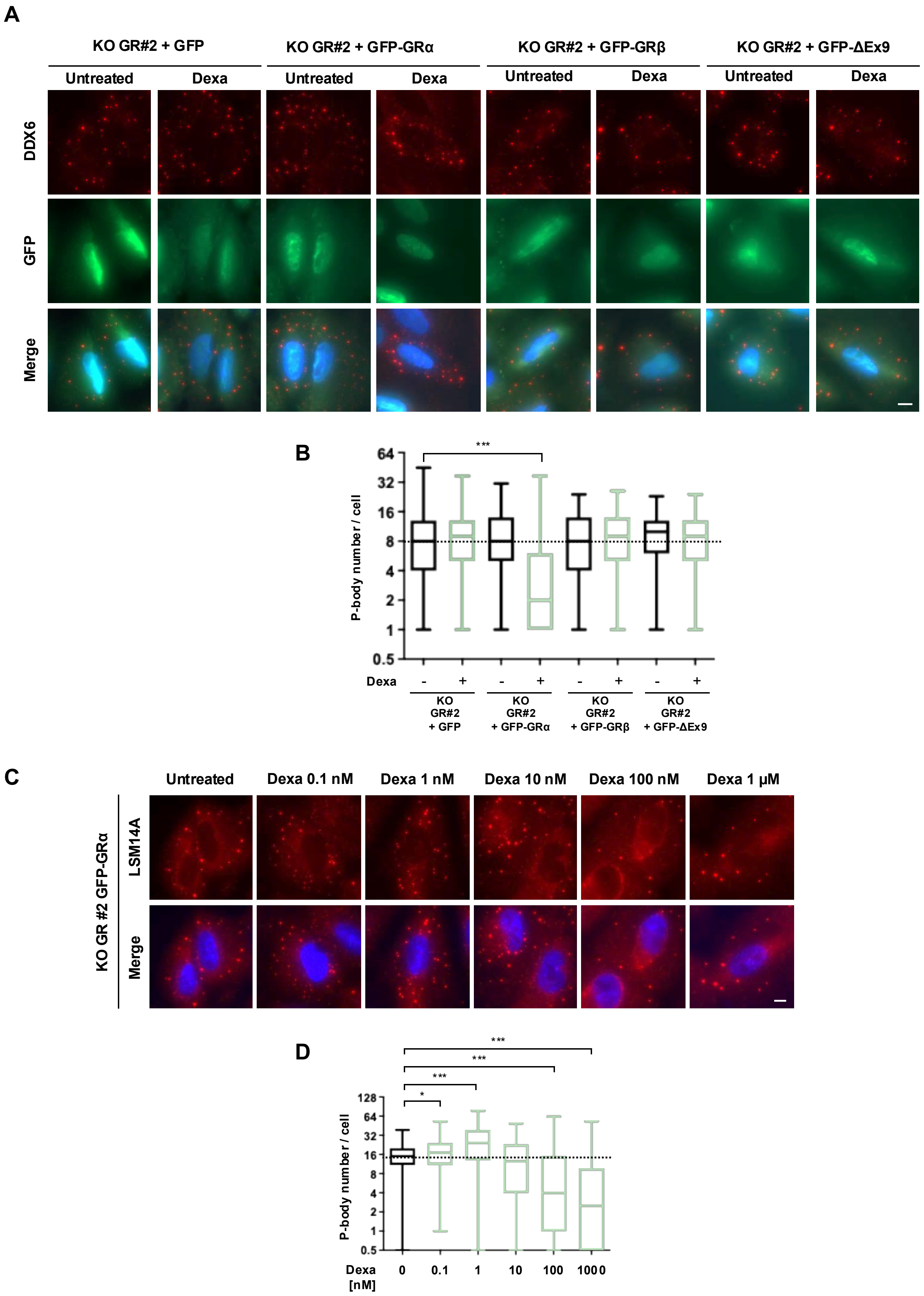


### Supp Figure 9

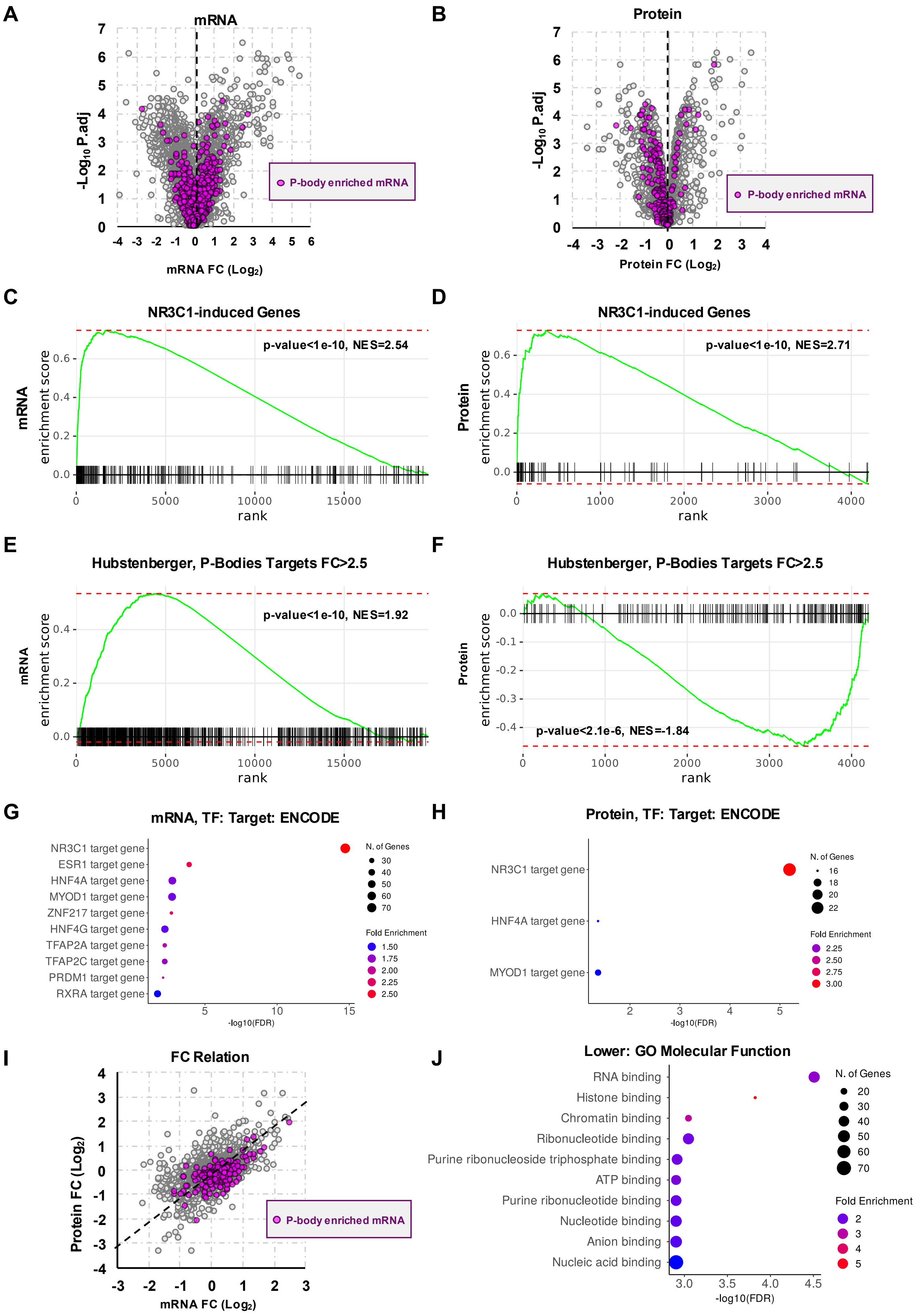


### Supp Figure 10

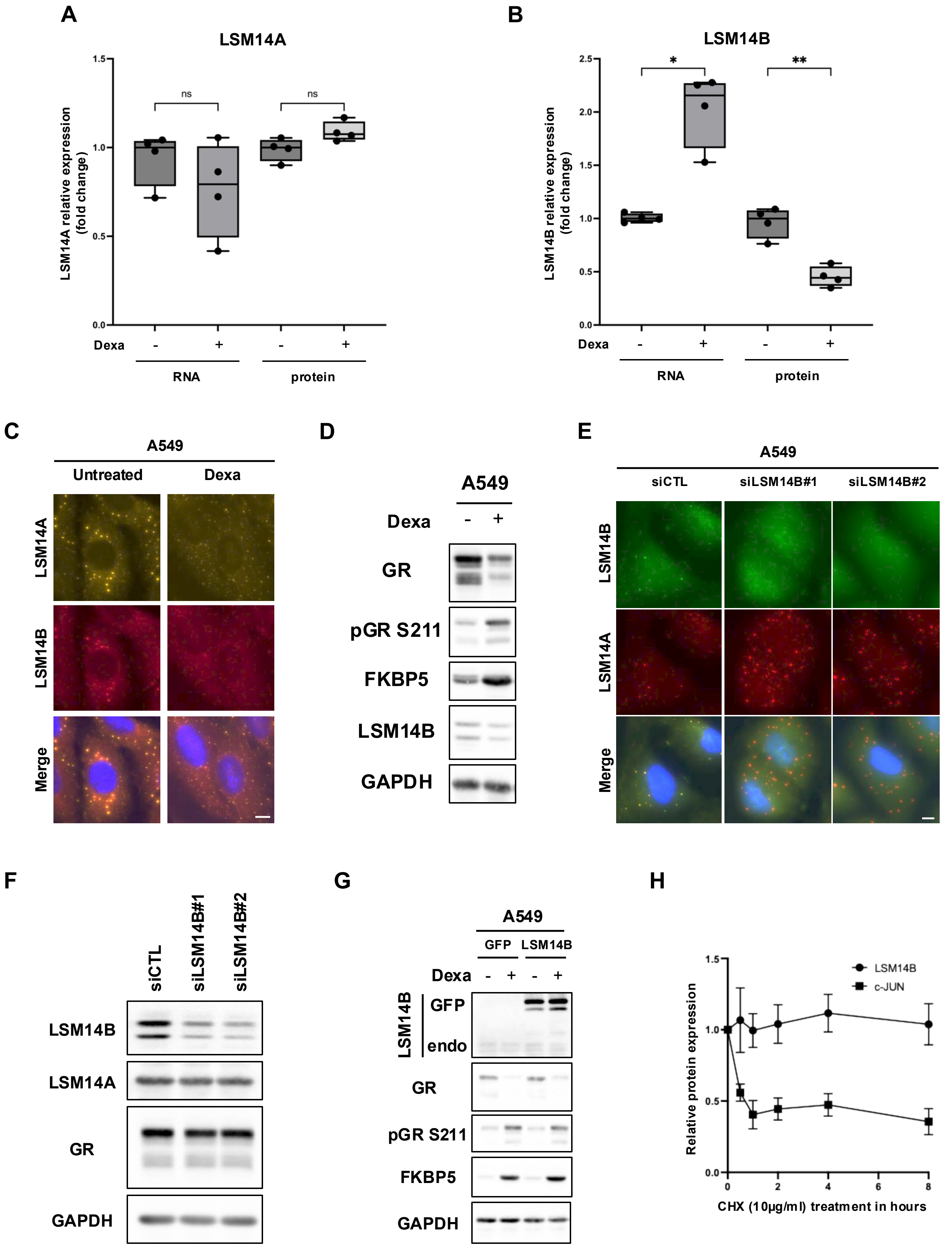


### Supp Figure 11

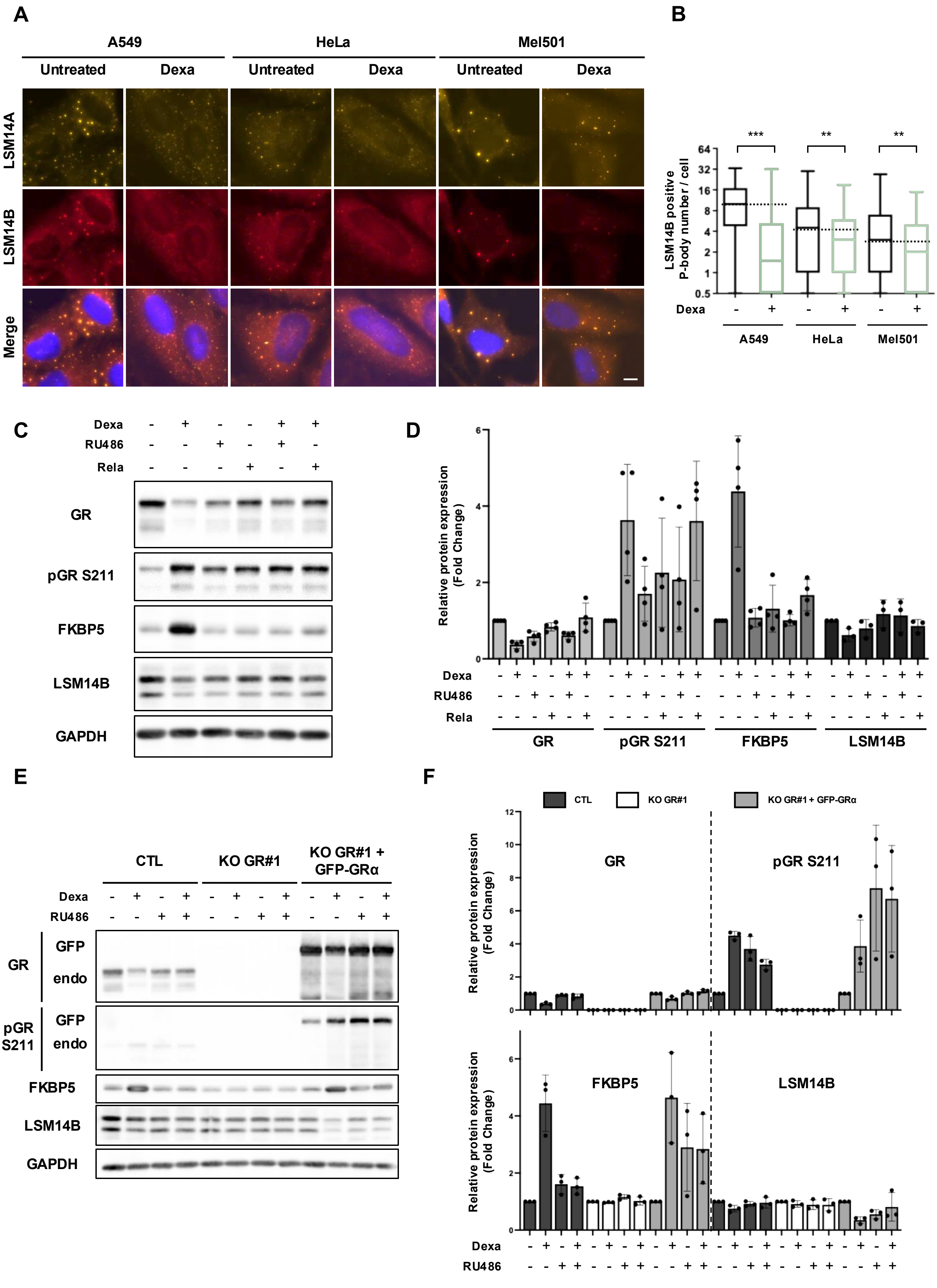


Supp Figure 12

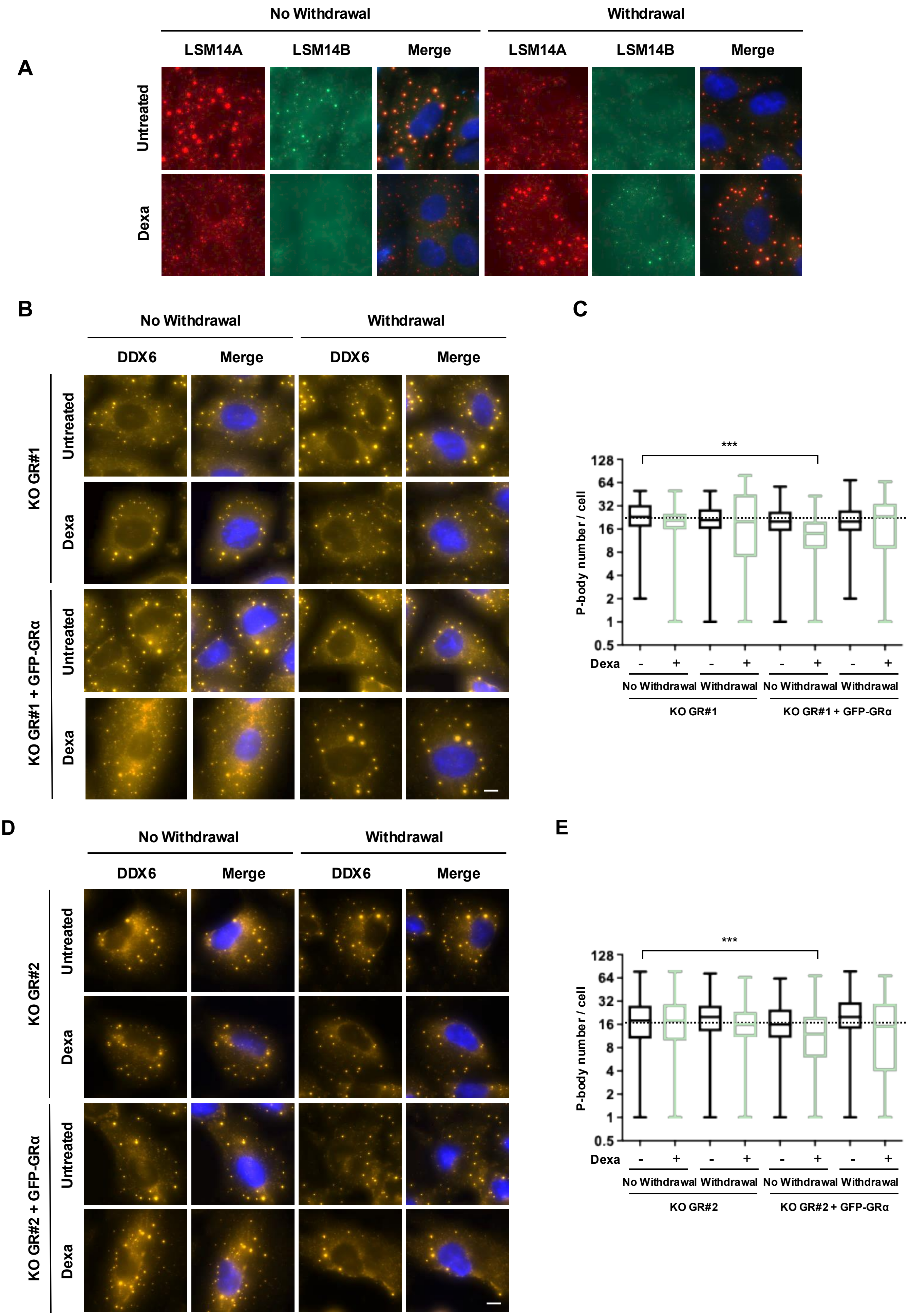


Supp Figure 13

### Graphical Abstract

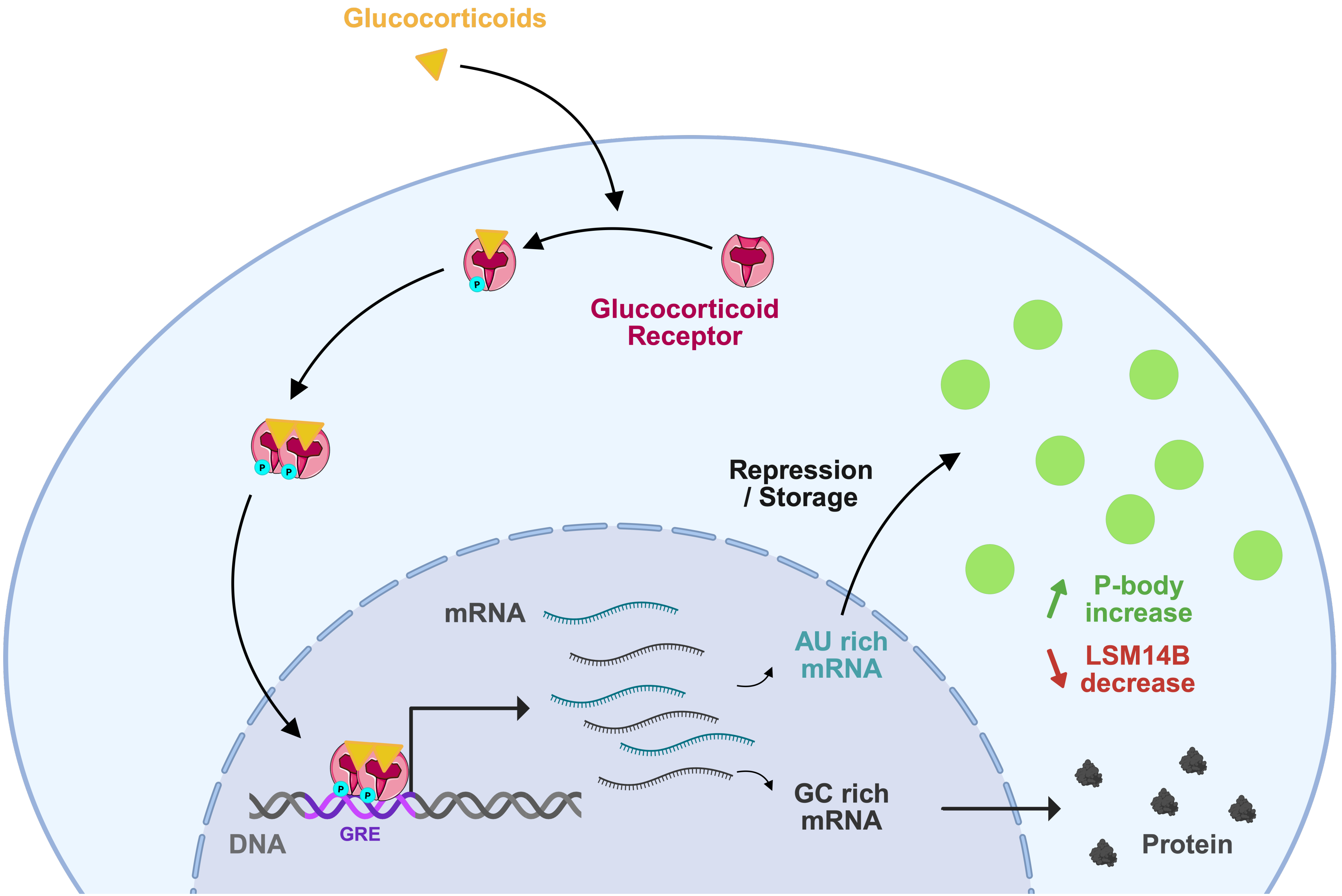
